# Intermediate Gray Matter Interneurons in the Lumbar Spinal Cord Play a Critical and Necessary Role in Coordinated Locomotion

**DOI:** 10.1101/2022.10.31.514612

**Authors:** Naëmi Kuehn, Andreas Schwarz, Carlo Antonio Beretta, Yvonne Schwarte, Francesca Schmitt, Melanie Motsch, Norbert Weidner, Radhika Puttagunta

## Abstract

Locomotion is a complex task involving excitatory and inhibitory circuitry in spinal gray matter. While genetic knockouts examine the function of unique spinal interneuron (SpIN) subtypes, the phenotype of combined premotor interneuron loss remains to be explored. We modified a kainic acid lesion to damage intermediate gray matter (laminae V-VII) in the lumbar spinal enlargement (spinal L2-L4) in female rats. A thorough, tailored behavioral evaluation revealed deficits in gross hindlimb function, skilled walking, coordination, balance and gait two-weeks post-injury. Using a Random Forest algorithm, we combined these behavioral assessments into a highly predictive binary classification system which strongly correlated with structural deficits in the rostro-caudal axis. Machine-learning quantification confirmed interneuronal damage to laminae V-VII in spinal L2-L4 correlates with hindlimb dysfunction. White matter damage and lower motoneuron loss did not correlate with behavioral deficits. Animals do not regain lost sensorimotor function three months after injury, indicating that natural recovery of the spinal cord cannot compensate for loss of laminae V-VII neurons. As spinal cord injuries are often located at spinal enlargements, this research lays the groundwork for new neuroregenerative therapies to replace these lost neuronal pools vital to sensorimotor function.

**Highlights:**

- Functional deficits in coordination, balance, rhythmic walking and gait follow two weeks after a lumbar (L2-L4) intermediate (V-VII) gray matter spinal cord injury in rats
- Deficits correlate with neuronal loss in laminae V-VII in spinal levels L2-L4 but do not correlate with lower motoneuron loss or white matter damage nor do animals show signs of sensory dysfunction due to spinal cord injury
- Coordination deficits remain after three months, indicating that natural recovery cannot compensate for this interneuronal loss
- Newly developed machine-learning models non-invasively fully classify injured animals by functional readouts equivalent to time-intensive endpoint histological analysis

## Introduction

There is a great deal of interest to better understand the role spinal interneurons (SpINs) play in motor function particularly in the cervical and lumbar enlargements (Dai et al., 2005; Guertin, 2009; Lanuza et al., 2004; Magnuson et al., 2005; Sathyamurthy et al., 2018; Zholudeva et al., 2021). Interconnected inhibitory and excitatory interneurons (INs) form plastic networks within these two spinal enlargements and are responsible for eliciting locomotor output and coordinating sensory and motor function for and between limbs. Gray matter neuronal loss particularly in these regions can lead to detrimental deficits (Hadi et al., 2000; Zholudeva et al., 2021).

It has been previously shown that a large gray matter lesion in the rostral-lumbar enlargement resulted in gross locomotor deficits. Gross hindlimb function was most severely affected when neuronal loss included spinal level L2; these deficits were not seen in a thoracic gray matter lesion highlighting the importance of neurons at this spinal level (Hadi et al., 2000; Magnuson et al., 1999). Furthermore, these deficits did not correlate with motoneuron loss or white matter damage. While it was suggested that the intermediate gray matter is responsible for these deficits, the most severe neuronal damage was seen in both the dorsal horns and intermediate gray matter (Hadi et al., 2000). Although damage to the dorsal horn is more commonly associated with sensory deficits, it is unclear whether this also played a role in behavioral deficits. Damage to the dorsal horn can evoke pain (Yezierski et al., 1998) and animals with pain in their hindlimbs can show altered gait patterns (Deshpande et al., 2021; Pitzer et al., 2016). Therefore, loss of afferent or other information due to the severity of the lesion may have impacted motor output.

Premotor IN ensembles in the intermediate gray matter receive and gate descending motor and ascending sensory input and elicit signals to other INs and motoneurons. Elegant cFos and genetic knockout/silencing experiments have revealed the functions of IN populations in laminae V-VII in the lumbar cord during locomotion in both mice and rats (Ahn et al., 2006; Flynn et al., 2011; Hayashi et al., 2018; Koch et al., 2017; Ruder et al., 2016; Sathyamurthy et al., 2018). While these experiments highlight the roles of individual SpIN subtypes, these studies have not shown the phenotype of combined premotor IN loss. Previous studies have shown that changes to the ratio of excitatory to inhibitory INs can influence the speed and pattern of the motor output as well (Sternfeld et al., 2017). For example, inhibitory V1 INs have been shown to be necessary for fast motor bursting. Ablation of V1 INs lengthened the step cycle and slowed motoneuron burst frequency (Gosgnach et al., 2006), while ablation of both V1 and V2b affects flexor-extensor alternation (Zhang et al., 2014). Furthermore, *in vitro* studies reconstructing SpIN circuits have shown that changing the ratio of excitatory and inhibitory neurons such as by adding more inhibitory V1 INs or excitatory V3 can influence rhythmic activity (Sternfeld et al., 2017). Therefore, changes to the regulatory SpIN network located in the intermediate gray matter, such as from a spinal cord injury, can affect motor output in an unpredictable manner. To our knowledge, it has not been shown what locomotor deficits are created after damage to the lumbar intermediate gray matter alone. With this study, we aim to build on the above-mentioned model and create a more specific gray matter lesion to thoroughly investigate the behavioral readout of premotor IN loss.

In this study, we examined the combined roles of INs and propriospinal INs in laminae V-VII in spinal levels L2-L4 in locomotion and whether intrinsic recovery after damage to this select region is possible. By modifying a spinal excitotoxic kainic acid (KA) lesion model (Magnuson et al., 1999), we were able to create a lesion targeting laminae V-VII including spinal levels L2-L4, lesioning both excitatory and inhibitory (Kerchner et al., 2001; Rodriguez-Moreno et al., 2000) local and propriospinal INs. We observed detailed acute locomotor deficits that were not recovered over a three-month period. We developed a novel Random Forest classification model to combine all behavioral assessments and segregate KA-lesioned from uninjured animals, allowing us to further compare performance and recovery over time. Machine-learning based neuronal quantification indicates that neuronal loss in spinal levels L2-L4 in laminae V-VII is critical and necessary for coordinated locomotion and cannot be compensated for by natural plasticity. This model allows us to further explore the potential of neurorestorative therapies, hence providing an ideal model for such studies.

## Material and Methods

### Experimental Design

#### Animals

Rats were housed in accordance with the European Union Directive and institutional guidelines. A total of 42 female Fischer 344 rats (Janvier Labs, Saint-Berthevin Cedex, France) weighing 180-200g (10 weeks old) at the time of the surgery were used for these experiments. 12 rats were used for pilot experiments with lesions in vertebrae T12. Animals with lesions that went into spinal level L2 were excluded from behavioral analysis (3 rats). Therefore, the total n-number was 9 rats, 4 controls and 5 KA animals. 30 rats were used vertebrae T13 injections. Of these, 21 rats were used in the first two-week experiment. One rat died after surgery and two control and 4 KA animals were excluded from the results as their lesion size did not fit the set criteria. Therefore, for the two-week experiment, a total of 14 rats were used, n = 7/group. For the three-month long-term experiment, a total of 9 rats were used. Two control animals and 1 KA animal were excluded for the same reason as mentioned above, resulting with a total of 6 rats, n=3/group. Rats were maintained in a 12hour light-dark cycle at 22 degrees Celsius. Rats were group-housed in Type IV cages, with a maximum of 6 rats per cage. Animals had ad libitum access to water and food throughout the experiment. All experiments were planned and conducted according to the PREPARE and ARRIVE guidelines (Percie du Sert et al., 2020; Smith, 2020).

#### Surgery

Experiments were conducted in accordance with the European Union Directive (Directive 2010/63/EU amended by Regulation (EU) 2019/1010) and institutional guidelines. Female Fischer 344 rats were injected intramuscularly with an anesthesia mix of xylazine, ketamine and acepromazine. After tail, paw and eye reflexes subsided, vertebra T12 or T13 was identified using the vertebra T10 landmark. For the pilot study, 12 animals received a laminectomy on vertebra T12 and received either one or two sets of bilateral 1µl, 1mM kainic acid (1mM; Tocris) injections. These injections were placed either rostral or rostral-mid vertebra T12. For the main experiment, 30 other animals received a laminectomy at vertebra T13 spanning 6mm. Following, three sets of bilateral 0.5µl injections of 1mM kainic acid were applied 0.4mm deep from the surface of the dura, 0.5mm laterally from the midline using a PicoSpritzer II (General Valve, Fairfield, NJ, USA) and a pulled glass capillary. Each set of injections was 3mm apart along the rostro-caudal axis. Control animals received saline injections at identical sites. Following, the overlaying muscles were sutured and skin was stapled. Animals were transferred to a warm heating pad until all reflexes had returned. All surgeries were performed by a blinded experimenter. Two days postoperatively, rats were given buprenorphine (analgesic) (0.03mg/kg; Reckitt Benckiser) and ampicillin (167 mg/kg; Ratiopharm) subcutaneously three times daily. Bladder function was also manually assessed twice per day during this time however rats did not display bladder dysfunction.

#### Behavioral testing

Behavioral testing was performed throughout the experiment to assess sensorimotor deficits or recovery. Only animals with weight support on both hindlimbs (BBB score ≥ 9) were tested on the horizontal ladder, inclined beam, von Frey, Hargreave’s and CatWalk. All behavior tests were performed by blinded examiners and were performed during the same time of day, in the morning, to maintain consistency.

#### Basso, Beattie, Bresnahan (BBB) Score and Subscore

For the two-week experiment, animals were tested 1, 3, 14 days after injury; for the three-month experiment, animals were tested 1, 3, 14, 30, 60 and 90 days after injury. Animals were placed into an open field where two blinded experimenters assessed gross hindlimb function (Basso et al., 1995). A subscore with a total of 13 points looked at hindlimb function dependent on coordination (including toe clearance, paw position, trunk stability and whether the tail was up or down).

#### Even and Uneven Horizontal Ladder

The horizontal ladder consisted of a custom, 1 meter setup with plexiglass and metal rungs. Rats were allowed to cross an elevated, unevenly spaced horizontal ladder (rungs spaced 2-3cm apart) five times to their home cage. After an hour break, animals were then allowed to cross an evenly spaced, elevated horizontal (rungs 3cm apart) ladder five times. Prior to baseline testing, animals were habituated on the evenly spaced ladder by allowing them to cross five times. Animals were not food deprived and no food reward was given at the end of the task. Following, animals in the two-week experiment were tested at baseline and 2 weeks after SCI. Animals in the three-month experiment were tested at baseline, 2, 4, 8, 12 weeks after injury. A Canon Legria HFR806 camera was placed perpendicular to the ladder set up and used to record performance. Animal fore- and hindlimb placement on the rungs were evaluated in slow motion using 25 frames per second and scored based on a previously established protocol (Metz & Whishaw, 2009). A new uneven rung pattern was tested at each time point to prevent a learning effect. Percent hindlimb slips and overall hindlimb performance scores were calculated and averaged.

#### Inclined Beam

Rats in the two-week experiment were tested on the inclined beam at baseline and 2 weeks after SCI. Rats in the three-month experiment were tested additionally 4, 8, and 12 weeks after injury. For this test, rats were placed onto an elevated and angled (10 degrees) 1.9cm narrow, wooden, circular 1 meter rod and trained to cross to reach their home cage. A green sheet was placed 0.3 meters below to gently catch rats who fell. Prior to baseline testing, animals were habituated by allowing them to cross five times or until they could cross without hesitation. For testing, rats were given the opportunity to cross the beam three times. A Canon Legria HFR806 camera was placed posterior to the beam and used to record inclined beam performance. Ability to complete the full task unassisted, time to cross and hindlimb slips were evaluated and scored. Each hindlimb step was given a maximum of 5 points, -1 point if foot placement was unstable; -2 points if the foot slipped off the beam but continued walking without stopping; -3 points the foot slipped off the beam and walking was interrupted; -4 points if the rat fell but was able to pull itself back up and complete the task; -5 points if the animal fell and did not complete the task. A time cut-off of 75 seconds was set. The total number of points was summed and normalized by the total possible number of points (total number of footsteps x 5). Scores for both hindlimbs over the three trials were averaged to calculate the average performance. Animals that did not complete the task received a 0.

#### Von Frey Filament Mechanical Sensitivity Testing

Dermatomes for the rat hindpaw lie in L4 and L5 (Takahashi et al., 2003), regions in which part of the KA lesion is located, allowing us to evaluate changes in sensitivity. During habituation, animals were placed in a small box on a metal grid (Ugo Basile) for one hour for three consecutive days. Following, mechanical sensitivity was tested at baseline, 1 and 2 weeks after injury in the two-week experiment. Animals in the three-month experiment were tested additionally at 4, 8, 12 weeks after injury. Von Frey filaments 1.4g, 4g, 8g, 16g, 60g were used to cover the spectrum of light touch to nociceptive stimuli. Positive responses on each hindpaw were recorded after five applications. There were 4 minutes between each consecutive stimulation to prevent desensitization. Hindpaw responses on both sides were averaged. Von Frey filament testing was only performed with animals that had weight support and full plantar steps for both hindlimbs.

#### Hargreave’s Thermal Testing

During habituation, animals were placed in a small box on a plastic floor (Ugo Basile) for one hour for three consecutive days. Following, baseline testing was performed in which a laser machine (50 units, maximum 30 seconds) was placed beneath each hindpaw and response time was recorded. Each hindpaw was tested four times and there were 8 minutes between each stimulation to prevent desensitization and damage. If there was urine or feces in the box, two minutes passed post-removal before testing resumed. This test was performed at baseline, 1, 2 weeks post-injury for animals in the two-week experiment. Animals in the three-month experiment were tested additionally 4, 8, 12 weeks post-injury.

#### CatWalk Gait Analysis

All animals were transported to the Interdisciplinary Neurobehavioral Core (INBC) in Heidelberg three days prior to testing for acclimatization. Following, a CatWalk Noldus XT was used to perform the CatWalk experiments. The following settings were used: camera height (48 cm); walkway width (7cm); green intensity threshold (0.1); camera gain dB (18.10); red ceiling light v (17.7); green ceiling light (16.0).

For this test, animals were trained to traverse the walkway without stopping. Habituation was performed one day prior to testing. On the day of the experiment, the first 5 runs where the maximum speed variation was under 50 and had speed variation less than ten were taken. All analysis was performed using the CatWalk Noldus XT software. The pLDA score was calculated according to the previously established protocol (Timotius et al., 2021). Animals without weight support were not included in the pLDA analysis. Following gait analysis, animals were transported back to the ZOUP Animal Facility Heidelberg for completion of the experiment. CatWalk gait analysis was performed once at the end of the two-week and three-month experiments, one day prior to sacrificing.

#### Classification of Animal Behavioral Performance

We were also interested in a correct segregation of KA-injured and non-injured animals without post-hoc assessment of the lesion-size. For this, extracted features were obtained by the performed behavioral tests (scores) and trained classification models using Random Forests which uses an ensemble of decision trees and aggregates their votes. Two independent approaches were explored: for the first approach (FULL), behavioral scores from all behavioral tests performed with the animals were extracted, namely (i) BBB, (ii) Horizontal Ladder, (iii) Inclined Beam, (iv) Sensory and the (v) CatWalk tests. For the second approach (ECO), the feature set was deliberately reduced to the BBB and inclined beam tests in order to provide a prediction model which relies only on a few fast and economically convenient behavioral tests.

Since the number of repetitions (trials) for most behavioral tests at each time point was different (1 to 5 repetitions), only 3 individual observations were generated per animal: For the Catwalk, horizontal ladder, inclined beam and all sensory tests from the first three trials were taken, for the BBB Scores the same score was used for each observation. In order to guarantee uniformity of the data, animals KA#5 and KA#7 were excluded from both classification approaches, since they did not perform the CatWalk test. In addition, all animals which were not able to successfully master the inclined beam test received 75s (stop criterion) for the time parameter, and 50 steps for the steps parameter. For each observation, 28 features were extracted: 2 BBB, 4 horizontal ladder, 4 inclined beam, 3 sensory and 15 CatWalk parameters. A detailed listing can be found in **Suppl.Tables 1 and 2**.

In this way, 21 observations were generated for the control group (3 × 7 control animals) and 15 for the KA group (3 × 5 KA animals), with a combined total of 36 observations (each observation included 28 behavioral parameters for the FULL model, and 6 behavioral parameters for ECO model).

Using a 10x 5 -fold cross-validation technique to avoid overfitting to the data, all available trials were divided into test and training data. A Random Forest classification model was trained to segregate observations between the control and KA group. One thousand trees were chosen for the decision aggregation and the square root of the number of features was chosen as the splitting criterion as previously published (Breiman, 2001). The mean of all aggregated votes and the accuracy obtained over all cross-validated results were reported. Chance was calculated using a Wald interval (aWI) for with a p value of 0.05 (Müller-Putz et al., 2008).

In addition, behavioral scores which contained the most discriminative information between KA-injured and non-injured animals were investigated. Using a Random Forest model, this feature ranking can be obtained by assessing the Gini Index for a feature at each split. The mean decrease of the Gini Index in magnitude was calculated for each feature over all cross-validated results.

Lastly, the votes obtained via decision aggregation per observation was assessed and correlated to the actual lesion size using Spearman’s correlation analysis.

### Tissue Processing and Immunohistochemistry

#### Transcardial Perfusion

All animals were transcardially perfused with saline and fixed with 4% paraformaldehyde (PFA). All cords were dehydrated in 30% sucrose for one week. Spinal cords were cryosectioned into 25 µm thick coronal sections in a 1:7 series on super frosted, charged glass slides. All tissue was stored at −20°C until use.

#### Immunohistochemistry

Slides were first dried for 15 minutes at room temperature. Slides were washed 3x with tris-buffered saline (TBS) and sections were permeabilized with a blocking solution of 5% donkey serum (Equitech-Bio Inc), 0.25% Tx-100 (neoLab Migge GmbH) in TBS for two hours at room temperature. Primary antibody (Merck guinea pig anti-NeuN 1:1000, Cat# ABN90) was added in 0.1% TX-100 and 1% donkey serum and incubated overnight at 4°C. The next day, slices were washed three times with 1% serum in TBS and incubated in secondary antibody (Dianova Alexa-Fluor 594 anti-guinea pig 1:300, Cat# 706295148) with DAPI (1:2000) for four hours at room temperature. Slides were washed again in TBS before being mounted on a cover-slip with Fluoromount G (Biozol, Cat # SBA-0100-01).

### Histological Analysis

#### Lesion Size and Exclusion Criteria

To determine the lesion size in the rostro-caudal axis, neuronal loss was identified and quantified in coronal sections immunostained for NeuN using an Olympus BX53 microscope. The number of slices with neuronal loss in a complete series were quantified and then multiplied by 25 microns x 7 to determine the length of the lesion. Within this region, the three slices with the greatest neuronal damage (ie the three bilateral injection sites) were identified as the lesion epicenters. All histological analysis was performed by a blinded experimenter. To determine if animals were appropriately lesioned and to ensure that the observed behavior is due to the desired lesion, we applied specific exclusion criteria based on the histological results. If control animals had a lesion size greater than 2800µm in the rostro-caudal extent, they were excluded from the experiment. If KA animals had a lesion size smaller than 6000µm, they were excluded from the experiment as they did not show consistent behavioral deficits.

#### NeuN Quantification in Laminae V-VII

To quantify NeuN-positive cells in laminae V-VII in an unbiased fashion, the following workflow was developed (**Suppl. Fig. 1**). Spinal cord sections were imaged using the Olympus XT1000 confocal microscope. Z-Stacks of 1.5 micron step size were acquired of each epicenter with the 10x magnification objective (UPlanSApo, 10x/0.40, infinity/0.17/FN26.5). Tiles were stitched in ImageJ/Fiji using the Grid/Collection stitching plugin with the following settings: Unknown Position; linear blending fusion method; 0.30 regression threshold; 2.5 max/avg displacement threshold; 3.5 absolute displacement threshold; and with subpixel accuracy (Preibisch et al., 2009). One coronal section (25µm) was analyzed per bilateral injection site (lesion epicenter). A total of three injection epicenters per spinal cord were analyzed.

To determine the correct spinal level, the maximum intensity projection (MIP) of the Z-stack of each injection epicenter was computed and registered with the matching spinal cord atlas section (Watson et al., 2009). A customized ImageJ/Fiji script was developed to semi-automate the registration process (https://github.com/cberri/2D_Registration_BrainAtlas_ImageJ-Fiji). The ImageJ/Fiji script uses the BUnwarpJ plugin (https://imagej.net/plugins/bunwarpj/) to register the MIP images with the corresponding selected atlas map (T13-L4). Indeed, the atlas section that best correspond to the spinal cord gray matter was selected and registered to the appropriate spinal section using land markers. The laminae V-VII regions of interest (ROIs) were cropped with ImageJ/Fiji polygon selection tool and saved (Schindelin et al., 2012); manual verification was performed. One coronal section (25µm) was analyzed per bilateral injection site (lesion epicenter). A total of three epicenters (6 ROIs) per spinal cord were analyzed.

To enhance signal to noise ratio and normalize the background across images, all the individual Z-stack tiles were processed using the ilastik pixel classification workflow (Berg et al., 2019). Two label classes (foreground NeuN and background) were used to differentiate NeuN-positive pixels from background pixels and the resulting foreground probability maps stitched in ImageJ/Fiji with the Grid/Collection stitching plugin. Ilastik pixel classification training was performed prior on 10 sample images. The 2D ROIs generated as described above were over-imposed on the 3D probability map and the pixels outside filled with zero values using the ImageJ/Fiji Clear Outside plugin. 3D automated image segmentation was performed on the processed ROI probability maps using cellpose *Nuclei* pretrained model in a customized Jupyter Notebook (cellpose version 0.6.2; (Stringer et al., 2021)). The labeled images were imported in arivis Vision4D (arivis AG) for visualization. Minor mistakes in the segmentation were manually corrected using arivis Vision4D 3D magic wand tool. The NeuN counts were exported in an Excel table and normalized by the selected ROI image volume (counts divided by (ROI area x 25microns)). Normalized NeuN counts for the respective left and right sides were summed to get the total NeuN count per volume at a specific spinal level. Total NeuN in L2-L4 was calculated by summing the total NeuN in each individual spinal level. To determine if there were differences between groups, the total averages per group were compared.

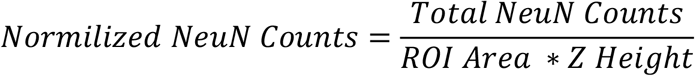

#### Workstation Hardware Specification

All image analysis was performed on a workstation equipped with a Nvidia RTX 2070 Super GPU, 128 GBs RAM and 10 Cores Intel i7 processor.

#### Software Accessibility

All custom code written for this project can be found under the following link: https://github.com/naemikuehn/lumbarcordanalysis.git

#### Motoneuron Analysis

To determine the number of remaining motoneurons at each lesion epicenter, NeuN-positive neuronal cell bodies in lamina IX were quantified using a similar method as previously published (Wen et al., 2015). The correct lamina was determine using the Spinal Cord Atlas (Watson et al., 2009). All NeuN-positive cells with a cell body larger than 919.632µm2 in laminae IX were counted and then analyzed in Graphpad Prism 6 to determine if there was a correlation between remaining motoneurons and behavioral performance. One coronal section was analyzed per bilateral injection site (lesion epicenter). A total of three epicenters per spinal cord were analyzed. All analysis was performed by a blinded investigator.

#### Percent white matter of total spinal cord cross-sectional area

To determine the percent white matter of the spinal cord cross-sectional area, a previously published protocol (Sliwinski et al., 2018) was used. Briefly, slides were subjected to an Eriochrome Cyanine staining and imaged using a XC30 camera and an Olympus Bx53 microscope (Olympus, Hamburg, Germany). Spared white matter was calculated in Image J by dividing the tissue sparing (total cross-sectional area of the slice – lesion size) and dividing it by the total cross-sectional area. One coronal section was analyzed per bilateral injection site (lesion epicenter). A total of three epicenters per spinal cord were analyzed. Percent white matter of the spinal cord was averaged per animal. An uninjured animal has 60-70% white matter. If an injury causes the gray matter to collapse, then the ratio between white and gray matter changes. All analysis was performed by a blinded investigator.

#### Heatmap Analysis

Behavioral tests and neuronal quantification at spinal levels L2-L4 were compared between individual KA-lesioned animals and the average controls. All values were normalized to the average controls and reported as a percentage of the average controls except for lesion size. For lesion size, all values were normalized to the largest lesion size, 15,925µm.

### Statistical Analysis

All data is shown as mean ± the standard error of the mean unless otherwise reported. A repeated measures 2-way ANOVA with Sidak’s post hoc multiple comparisons test was calculated to evaluate differences between the two groups in tests that were repeatedly performed over a period of time (BBB score, BBB subscore). Unless otherwise noted, all reported p-values are group differences p-values. All other tests underwent Shaprio-Wilk’s normality test. If normally distributed, a Welch’s unpaired t-test to determine differences between the two groups, if not normally distributed, a Mann-Whitney test was used (inclined beam, horizontal ladder, von Frey Filament testing, Hargreave’s, CatWalk, motoneuron, white matter sparing, NeuN counts). Linear regression analysis was performed to compare lesion size, motoneuron counts and white matter sparing with behavior. Shapiro-Wilk’s normality test was performed prior to Pearson’s (normally distributed) or Spearman’s (not normally distributed) correlation analysis to analyze the relationship between the two variables. All statistical tests were performed with Prism 6 and Prism 9 software (Graphpad, San Diego, CA, USA).

### Schematics

Schematics were created with BioRender.com, powerpoint and google images.

## Results

### Targeting intermediate gray matter for neuronal loss in lumbar spinal levels L2-L4 with three sets of bilateral KA injections

In order to create a lesion that targeted various SpINs contributing to locomotion, KA injection parameters were refined to create selective intermediate gray matter damage in the lumbar cord with a behavioral readout. This was achieved with three bilateral KA injections performed at vertebral level T13 in female Fischer rats, each consisting of 0.5µl of 1mM KA injected 0.5mm from the midline at a depth of 0.4mm (**Fig. 1*A-C***). All gray matter neurons were visualized with NeuN with a custom workflow to confirm damage in the target region. The ImageJ/FIJI plugin “registration of pairs” was used to align the spinal atlas over 3D confocal coronal section Z-stacks for determination of the spinal level and target laminae (**Fig. 1*D***). To quantify the number of NeuN-positive neurons in laminae V-VII from spinal levels L2-L4 in a consistent and unbiased fashion, the newly developed combined image analysis workflow was used: ilastik pixel classification, 3D segmentation using cellpose and arivis Vision4D for visualization and segmentation correction (**Suppl. Fig. 1**). Kainate lesions significantly reduced the number of NeuN-positive neurons in laminae V-VII to 14.39 ± 1.92 neurons/µm3*10-5 in comparison to 24.91 ± 1.89 neurons/µm3*10-5 in the control group (**Fig. 1*E***). With this experiment, we have confirmed that injecting KA with these defined parameters creates intermediate gray matter damage in the spinal levels L2-L4 of the spinal cord.

**Figure 1.**
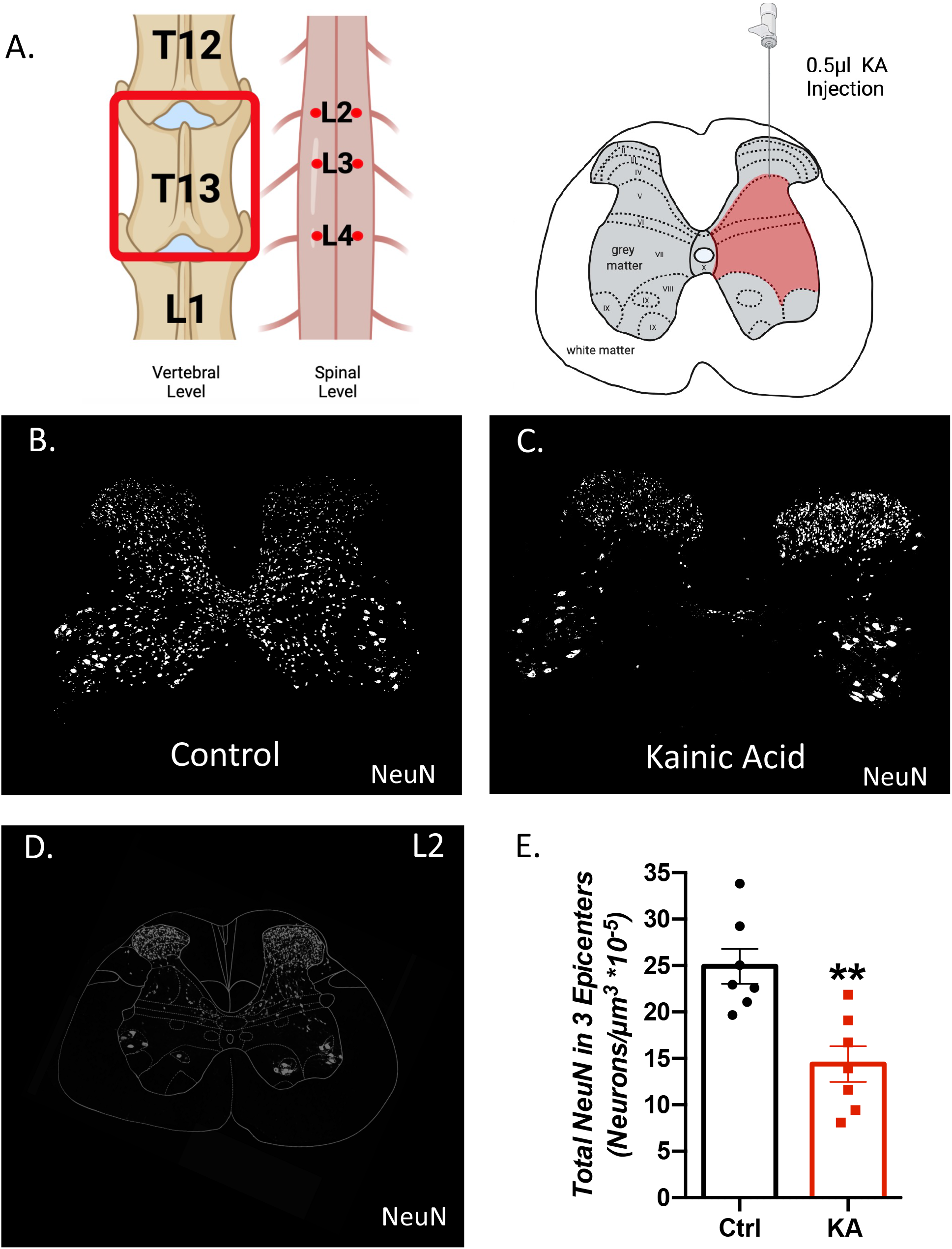
Defined kainic acid (KA) injection parameters creates selective intermediate gray matter damage in the lumbar spinal cord. ***A***, Schematic shows vertebral and spinal injection levels in the rostral and caudal axis as well as in the dorsal-ventral axis. ***B***,***C*** Representative images of lesion epicenters stained with NeuN to visualize neuronal loss in the control and kainic acid spinal cord. ***D***, Representative image of atlas overlay on coronal spinal cord section at spinal L2. ***E***, Total NeuN in laminae V-VII at lesion epicenters were quantified and compared (Welch’s unpaired t-test, ** p = 0.0021, n = 7 animals per group).

### KA rats show deficits in gross hindlimb function, rhythmic and skilled walking, coordination, balance and gait after two weeks

To assess the behavioral deficits induced by a discrete KA gray matter SCI, we performed a series of behavioral tests two weeks post-injury. The Basso-Beattie-Bresnahan (BBB) was performed 1, 3, 7 and 14 days post-injury (dpi) to evaluate gross hindlimb motor function (**Fig. 2*A,B***). We observed animals with a KA lesion have significant deficits in gross hindlimb function 1 day after injury which stabilizes one-week post-SCI (**Fig. 2*A***: Mean BBB scores after 14 days: control = 17.57 ± 0.69, KA = 11.21 ± 2.25) as well significant deficits in hindlimb function dependent on coordination (**Fig. 2*B***: mean BBB subscore control = 11.29 ± 0.18, KA = 4.57 ± 1.67). These deficits correlate to lesion size in the rostro-caudal axis (**Fig. 2*C,D***: BBB score r = −0.8254 and p= 0.0004, BBB subscore r = −0.7138 and p = 0.0041, Spearman’s correlation).

**Figure 2.**
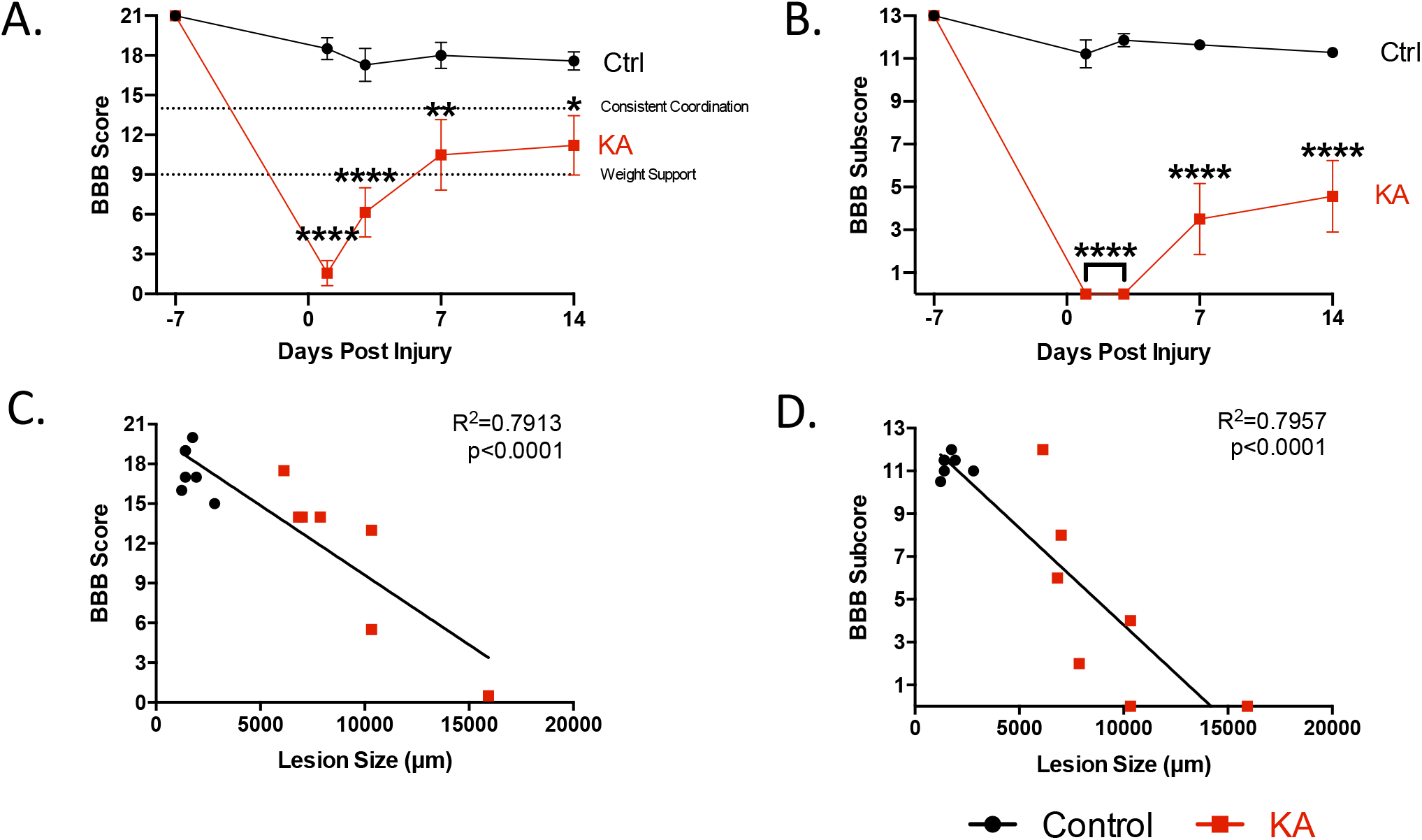
KA-injected rats display deficits in gross hindlimb function after two weeks. A score below 9 indicates that animals do not have full weight-supported steps but a score above 14 indicates consistent coordination while walking. ***A***,***B*** KA-injected animals have a significantly lower BBB score and sub-score after two weeks in comparison to the control (BBB score: 2-way ANOVA with Sidak’s post hoc test, group p < 0.0001; BBB sub-score: 2-way ANOVA with Sidak’s post hoc test, group p < 0.0001). ***C***,***D*** There were significant correlations between lesion size in the rostro-caudal axis and BBB score and sub-score (Linear Regression Analysis, BBB score p < 0.0001, R^2^ = 0.7913; BBB sub-score p < 0.0001, R^2^ = 0.7957). For all data, n = 7 animals per group; * p ≤ 0.05, ** p ≤ 0.01, **** p ≤ 0.0001.

Although the BBB score is useful in characterizing SCI animals, it is not as sensitive for tasks we believe are affected by our lesion model. Therefore, we opted for tasks specific for our lesion paradigm such as rhythmic and skilled walking, where animals were tested on an even horizontal ladder, uneven horizontal ladder (Martins et al., 2022) and inclined beam (Carter et al., 2001) (**Fig. 3*A,D,G***). KA rats show significantly higher number of hindlimb slips on both the even and uneven horizontal ladders indicating deficits in rhythmic walking (**Fig. 3*B***: even ladder controls: 0.70 ± 0.34%, KA = 39.91 ± 15.81%) and coordination (**Fig. 3*E***: uneven ladder controls: 1.71 ± 0.91%, KA = 48.68 ± 15.12%). Only 3 of the 7 KA rats were able to complete the inclined beam test (**Fig. 3*H***: controls = 100 ± 0% completion, KA = 23.81 ± 14.02%). Those KA rats which did complete the beam had a lower performance score (Controls: 95.66 ± 1.25%, KA: 21.19 ± 13.32%) further indicating deficiencies in coordination and balance (**Suppl. Video**). These behavioral tests significantly correlated to lesion size in the rostro-caudal axis (**Fig. 3*C,F,I***: even horizontal ladder r = 0.8490 and p = 0.0003, uneven horizontal ladder r = 0.8401 and p = 0.0004, inclined beam r = −0.8944 and p < 0.0001, Spearman’s correlation).

**Figure 3.**
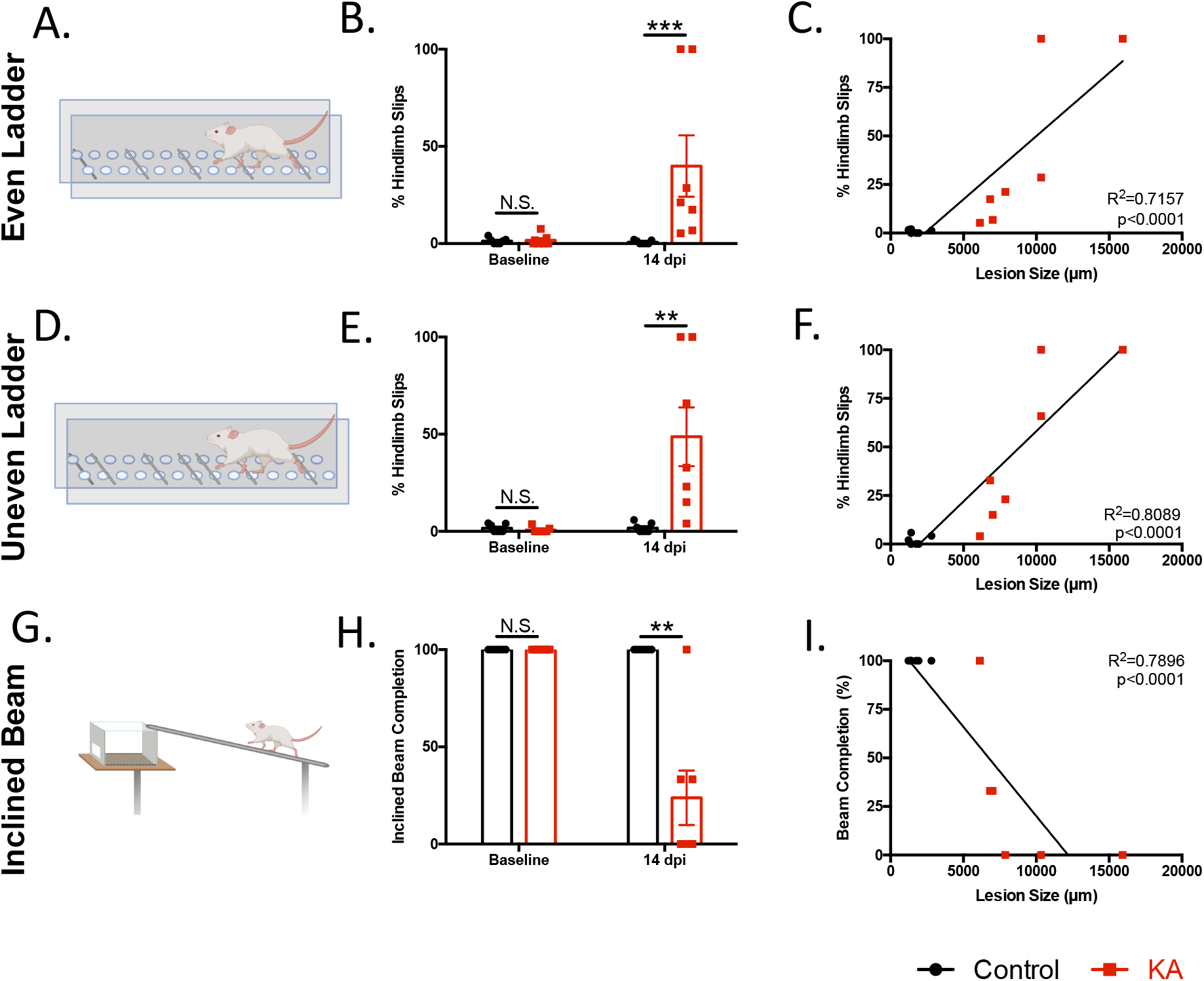
KA-injected rats display deficits in rhythmic and skilled-walking, coordination and balance, correlating to lesion size. ***A***, Schematic of the evenly-spaced rungs of the horizontal ladder. ***B***, Percent hindlimb slips were compared two weeks post-KA SCI (Mann-Whitney test, at baseline p = 0.9184, at 14 dpi p = 0.0006). ***C***, Correlation analysis is significant between hindlimb slips and lesion size in the rostro-caudal axis (Linear Regression Analysis, p < 0.0001, R^2^ = 0.7157). ***D***, Schematic of the unevenly spaced rungs of the horizontal ladder. ***E***, Percent hindlimb slips were compared two weeks post-KA SCI (Mann-Whitney test, at baseline p = 0.4126, at 14 dpi p = 0.0023). ***F***, Correlation analysis is significant between hindlimb slips and lesion size in the rostro-caudal axis (Linear Regression Analysis, p < 0.0001, R^2^ = 0.8089). ***G***, Schematic of the inclined beam behavioral test. ***H***, Ability to complete the inclined beam task was significantly impaired in KA SCI rats (Mann-Whitney test, at 14dpi p = 0.0047) ***I***, Correlation analysis is significant between ability to complete the inclined beam and lesion size in the rostro-caudal axis (Linear Regression Analysis, p < 0.0001, R^2^ = 0.7896). For all data, n = 7 animals per group; N.S. stands for not significant, ** p ≤ 0.01, *** p ≤ 0.001.

Static and dynamic parameters of each forelimb and hindlimb as well as the whole-body automated gait analysis was performed with the CatWalk (Noldus). This test also further investigates the role of propriospinal INs connecting the cervical and lumbar enlargements. An algorithm (parameter-combined linear discriminant analysis, pLDA) that combines 9 SCI-related gait parameters found to be predictive of injured vs uninjured thoracic SCI rat models was applied. In comparison to the controls, KA rats have a significantly lower pLDA score two weeks after SCI (**Fig. 4*B***: controls = 0.13 ± 0.01, KA = 0.076 ± 0.01). Separation into the nine individual parameters indicated significant differences in all except for hindlimb base of support and alternate B (AB) step sequence (**Fig. 4*D*)**. Additionally, we found KA-lesioned rats had a significantly higher cruciate A (CA) step sequence in comparison to the controls (**Fig. 4*A,C*:** controls = 10.48 ± 4.50%, KA = 27.41 ± 5.46%). A closer analysis of the rhythmic components of gait showed significant differences in forelimb swing time, stand time and duty cycle **(Suppl. Fig. 2***A,C,E*). However, the hindlimb parameters were not significantly different (**Suppl. Fig. 2***B,D,F)*.

**Figure 4.**
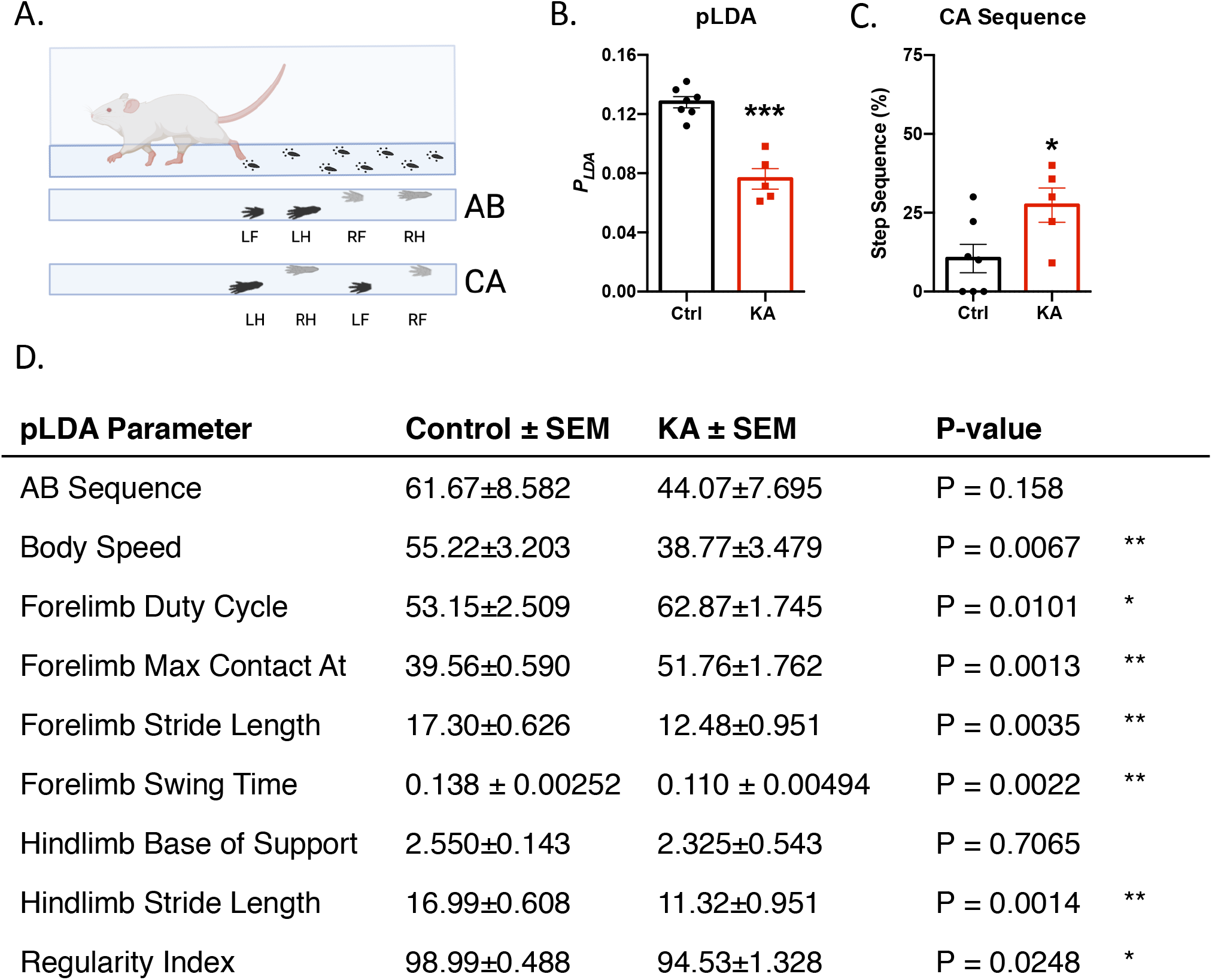
CatWalk gait analysis highlights deficits in SCI-related gait parameters two weeks post-injury. ***A***, Schematic of the CatWalk behavioral test with AB (alternate) vs CA (cruciate) step sequences. The AB step sequence requires a shift from hind-to forelimb (or fore-to hindlimb) on the ipsilateral side of the body while the CA step sequence requires a shift from fore-to forelimb (or hind-to hindlimb) on the contralateral side. ***B***, Comparison of the pLDA gait score shows significant differences (Welch’s unpaired t-test, p = 0.0006). ***C***, Comparison of CA sequence shows significant differences (Welch’s unpaired t-test, p = 0.0415). ***D***, Analysis of individual pLDA CatWalk parameters. Significance determined using Welch’s unpaired t-test. For all data n = 7 control and n = 5 KA animals; * p ≤ 0.05, ** p ≤ 0.01, *** p ≤ 0.001.

To further investigate the specificity of our lesion and ensure we did not disrupt the dorsal horn circuitry that may lead to sensory deficits (Vierck et al., 2013; Yezierski et al., 1998), we performed several sensory tests. We did not find any significant differences in mechanical and thermal sensitivity in animals two weeks post-KA SCI (**Fig. 5*A-C***). These results suggest that KA did not damage the dorsal horn as there were no significant sensory differences after two weeks.

**Figure 5.**
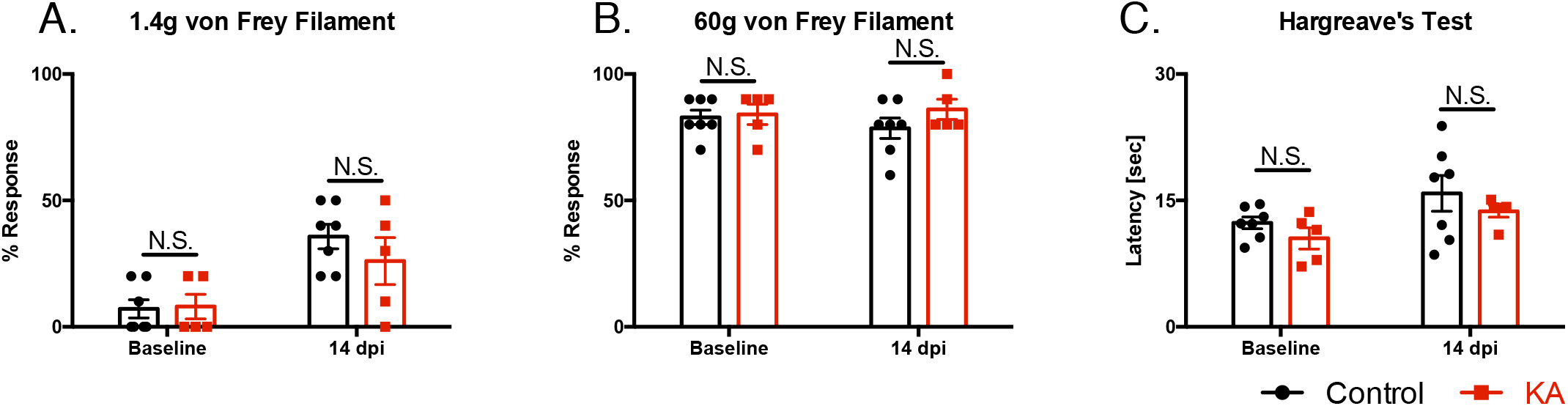
KA animals do not show differences in mechanical or thermal sensitivity after two weeks. ***A***,***B*** Hindpaw stimulation with 1.4g innocuous and 60g noxious von Frey hair filaments did not show significant differences at baseline or after two weeks (Mann-Whitney test, 1.4g at baseline p > 0.9999; Welch’s unpaired t-test, 1.4g at 14 dpi p = 0.3875; 60g at baseline p = 0.8222, 60g 14 dpi at p = 0.2217). ***C***, Hargreave’s hindpaw stimulation did not show differences in thermal sensitivity between the two groups (Welch’s unpaired t-test, at baseline p = 0.2496, at 14 dpi p = 0.3727). For all data, n = 7 control and n = 5 KA animals. N.S. stands for not significant.

Together, this battery of behavioral assessments of rats with a lumbar intermediate gray matter SCI shows gross hindlimb deficits, impaired rhythmic walking, coordination, balance and gait but no differences in sensory function two weeks post-injury. Lesions in a mixed cohort which we found to be primarily in spinal levels T13/L1 and not overlapping into spinal L2 did not show significant gross hindlimb and coordination deficits at baseline and two weeks after injury. At baseline the BBB score of both the control and KA groups was 21 ± 0.0. At 14dpi, the BBB score of the control group was 20.63 ± 0.38 and the KA group was 18.20 ± 0.92 and not significantly different (p = 0.0642, unpaired Welch’s t-test, n = 4 control and n = 5 KA animals). The BBB subscore at baseline for both the control and KA groups was 13.00 ± 0.0. Two weeks after injury, the BBB subscore for the controls was 12.75 ± 0.25 and for the KA animals was 10.8 ± 0.80 and was also not significantly different (p = 0.0750; unpaired Welch’s t-test, n = 4 control and n = 5 KA animals). Finally, the percent hindlimb slips on the uneven ladder for the controls at baseline was 2.15 ± 1.44 and for the KA animals was 2.26 ± 1.20% (p = 0.9557, unpaired Welch’s t-test, n = 4 control and n = 5 KA animals). Two weeks post lesioning, the percent hindlimb slips for the controls was 1.19 ± 0.79% and for the KA group was 2.78 ± 1.15% (p = 0.3194, unpaired Welch’s t-test, n = 4 controls, n = 5 KA animals) Together, this indicates that the observed deficits are specific to this lesion in spinal L2-L4.

### Neuronal quantification highlights role of spinal level L2-L4 INs in coordination and balance

We next wanted to determine whether neuronal loss at a specific spinal level significantly contributes to behavioral deficits. Using the above-mentioned NeuN quantification method in Figure 1, we further analyzed the number of NeuN-positive neurons in laminae V-VII in spinal L2-L4 in KA animals and normalized this to the average controls. In comparison to the average controls, there is a reduction of NeuN in all KA injected spinal levels except for KA#1 in spinal level L2 (**Fig. 6*A***). We found that the number of NeuN-positive neurons in all three spinal levels L2-L4 significantly correlates to the inclined beam performance (**Fig. 6*B-D***: L2 p = 0.0187 and r = 0.6325; L3: p = 0.0219 and r = 0.6371; L4: p = 0.0021 and r = 0.8029, Spearman’s correlation, respectively). This data indicates that intermediate gray matter IN loss in all three spinal levels (L2-L4) correlates with coordination and balance performance **(Suppl. Fig. 3)**.

**Figure 6.**
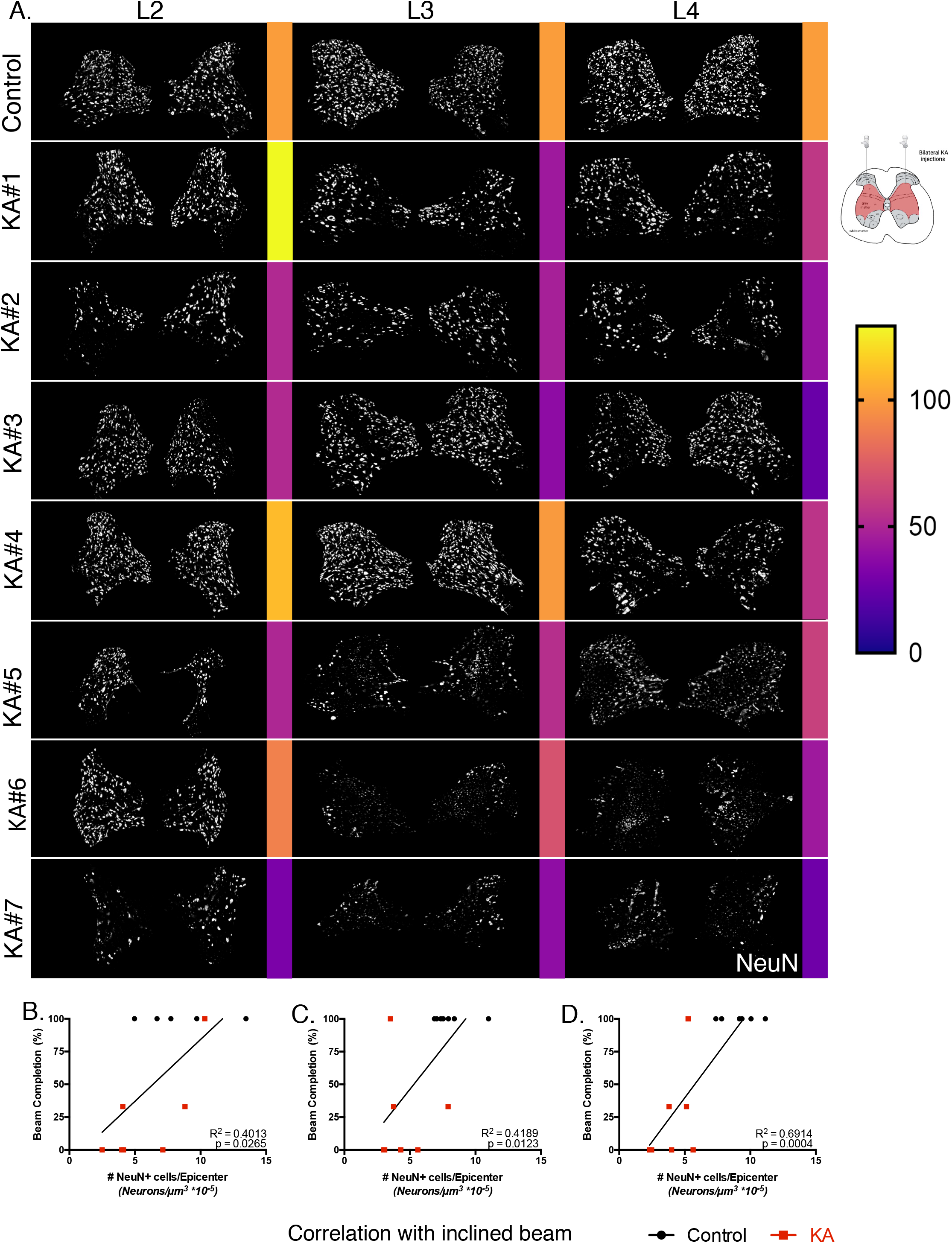
Laminae V-VII neuronal loss in spinal levels L2-L4 correlates with deficits in balance and coordination deficits. ***A***, Visualization of NeuN in laminae V-VII in spinal levels L2-L4 using ilastik MIPs. Single images of the KA animals in comparison to a representative control animal. Number of neurons are quantified and normalized to the average of the controls, shown as a percentage (0% in violet, 100% in orange, above 100% in yellow). ***B-D***, Correlation of NeuN in spinal level L2-L4 in comparison to inclined beam performance (Linear Regression Analysis, L2 inclined beam p = 0.0265, R^2^ = 0.4013; L3 inclined beam p = 0.0123, R^2^ = 0.4189; L4 inclined beam p = 0.0004, R^2^ = 0.6914; n = 5-7 animals per group).

### Lower motoneurons and white matter tracts are not damaged by spinal L2-L4 intermediate gray matter KA-SCI

Having shown that neuronal loss in laminae V-VII correlates with the observed deficits, we wanted to investigate if there was any correlation between motoneuron or white matter loss and behavior. To examine if motoneurons were targeted by KA, we counted the number of lower motoneurons at the injection epicenters and correlated them to the BBB score and inclined beam, and found no significant correlations (**Fig. 7*A,B***: BBB r = −0.2327 and p = 0.388, inclined beam r = −0.2338 and p = 0.226, Spearman’s correlation).

Although KA targets gray matter primarily, we wanted to ensure we did not secondarily damage the white matter tracts, therefore we examined the percent white matter of the spinal cord cross-sectional area by eriochrome cyanine staining. No significant correlation between percent white matter of the cord and BBB score or inclined beam was found (**Fig. 7*C,D***: BBB r = −0.1333 and p = 0.664; inclined beam r = −0.4221 and p = 0.103, Spearman’s correlation). Taken together, these results show that motoneuron loss and white matter damage did not contribute to the observed behavioral deficits.

**Figure 7.**
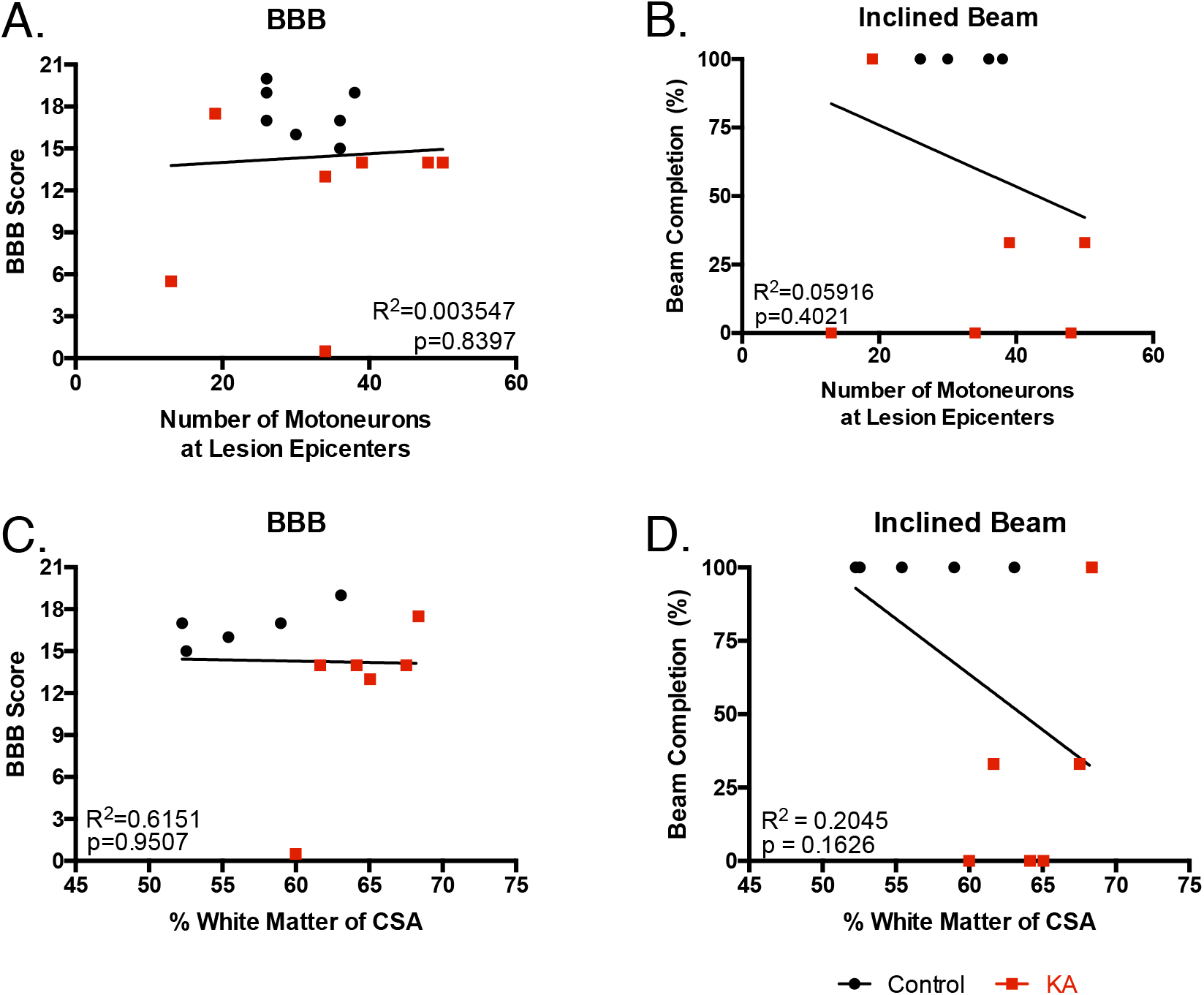
Behavioral deficits do not correlate with white matter damage nor motoneuron loss at spinal levels L2-L4. ***A, B*** Sum of remaining motoneurons at lesion epicenters does not correlate with BBB score nor inclined beam completion (Linear Regression Analysis, BBB score p = 0.8397, R^2^ = 0.003547; Beam Completion p = 0.4021, R^2^ = 0.05916, n = 7 animals per group). ***C, D*** Percent white matter of the cord cross sectional area (CSA) does not correlate with behavioral tests (Linear Regression Analysis, BBB score p = 0.9507, R^2^ = 0.6151; Inclined beam p = 0.1626, R^2^ = 0.2045, n = 5 control and n = 6 KA animals).

### KA lesioned animals do not recover over time

Endogenous plasticity is thought to help regain function after thoracic SCI and should be evaluated over time. KA-lesioned rat behavioral performance three months post-injury reveal that deficits in gross hindlimb function and coordination remain (**Fig. 8*A,B***, **Suppl. Fig. 4**). A closer look at the BBB score and subscore show that KA-injured rats have significant deficits in hindlimb function (**Fig. 8*A,B***: BBB score controls = 19.83 ± 0.60, KA = 10 ± 2.5; BBB subscore controls = 12 ± 0.58, KA = 2.33 ± 2.33). Two of the three animals did not regain weight support and were not able to participate in the ladder and beam tasks which increases the variability observed (**Suppl. Fig. 4**). However, from the BBB score and subscore it appears that the KA lesioned animals did not recover. We found that while lesion size did correlate with the gross hindlimb function (**Fig. 8*C,D***: BBB score r = −0.8971, p = 0.0110, BBB subscore r = −0.8971, p = 0.0111, Spearman’s correlation), the motoneuron number at the injection epicenters did not (**Fig. 8*E,F***: BBB score r = −0.1160, p = 0.7778; BBB subscore r = −0.1160, p = 0.7778, Spearman’s correlation). This suggests that the targeting of the intermediate gray matter of the L2-L4 spinal cord leads to permanent locomotor deficits.

**Figure 8.**
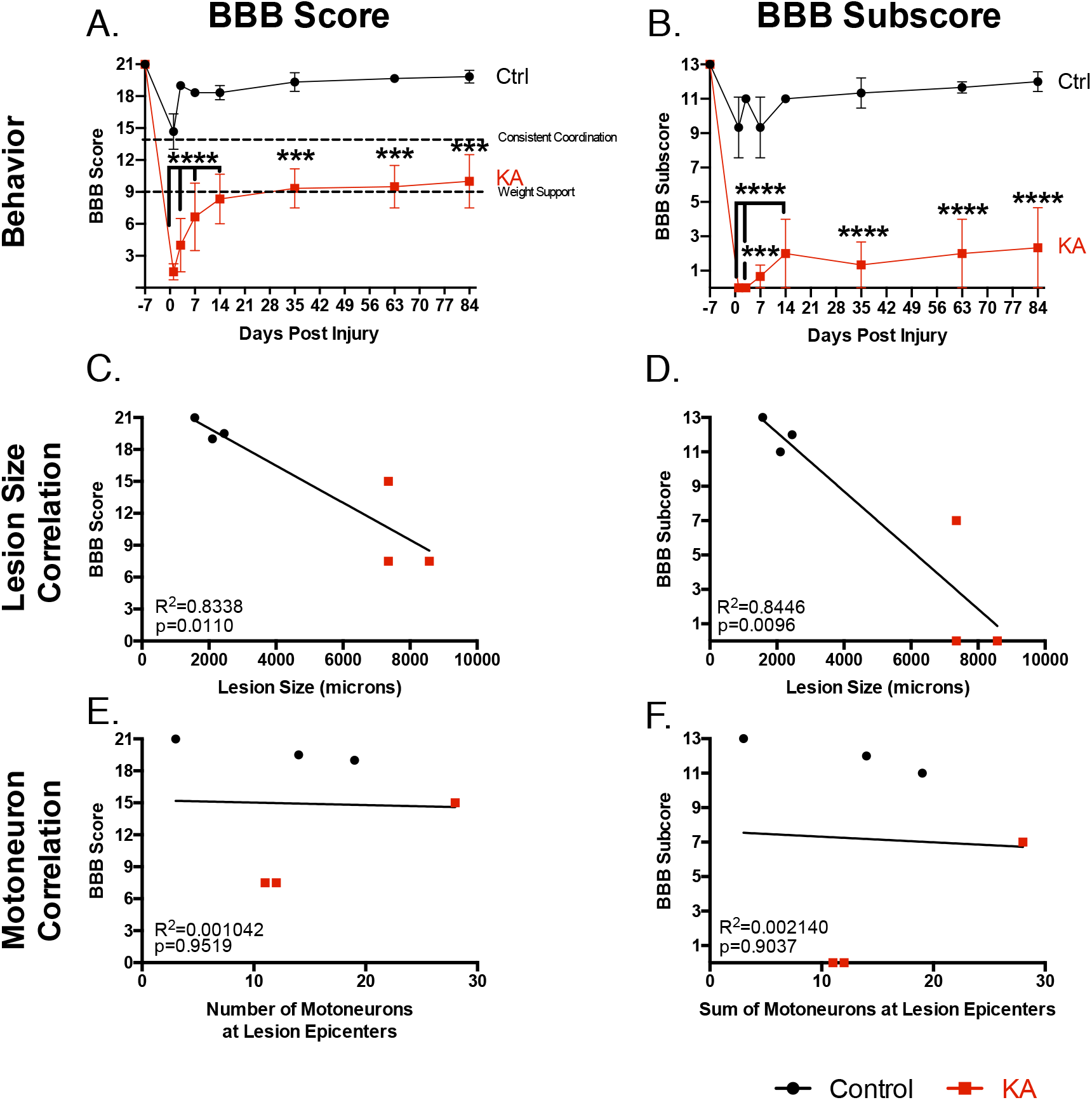
Analyzing the behavioral stability of the lesion after three months shows gross hindlimb and coordination deficits remain. ***A***,***B*** BBB score and subscore over a three-month period shows that gross hindlimb functional deficits remain (2-way ANOVA with Sidak’s post hoc test, BBB group p = 0.0062; BBB subscore group p = 0.0013). ***C, D*** BBB score and subscore correlate significantly to lesion size after three months (Linear Regression Analysis, BBB score p = 0.0110, R^2^ = 0.8338; BBB subscore p = 0.0096, R^2^ = 0.8446). ***E, F*** Remaining motoneurons at lesion epicenters do not correlate to behavioral performance after three months (Linear Regression Analysis BBB score p = 0.9516, R^2^ = 0.001042). For all data, n = 3 animals per group; *** p ≤ 0.001, **** p ≤ 0.0001.

### Comprehensive behavioral classification

Each behavioral assessment provides unique information on deficits induced by KA injury. However, performance variation made it difficult to classify animals prior to post-mortem histological analysis. Using a Random Forest classification approach, we were able to develop two models that classify animals into their appropriate test groups (**Fig. 9**). For the comprehensive and sensitive FULL model, all available behavioral parameters mentioned above (BBB, horizontal ladder, inclined beam, sensory and CatWalk tests) were used with a mean accuracy of 94.8 ± SD 8.2% (chance level 63%, p<0.05, adjusted Wald Interval) reached. The confusion matrix indicates that 12.7% of the KA observations were misclassified as control (Type 2 error). The mean of the normalized votes indicates that 11 out of 12 animals were appropriately classified animals into either the control or KA groups (**Fig. 9*A,C***).

**Figure 9.**
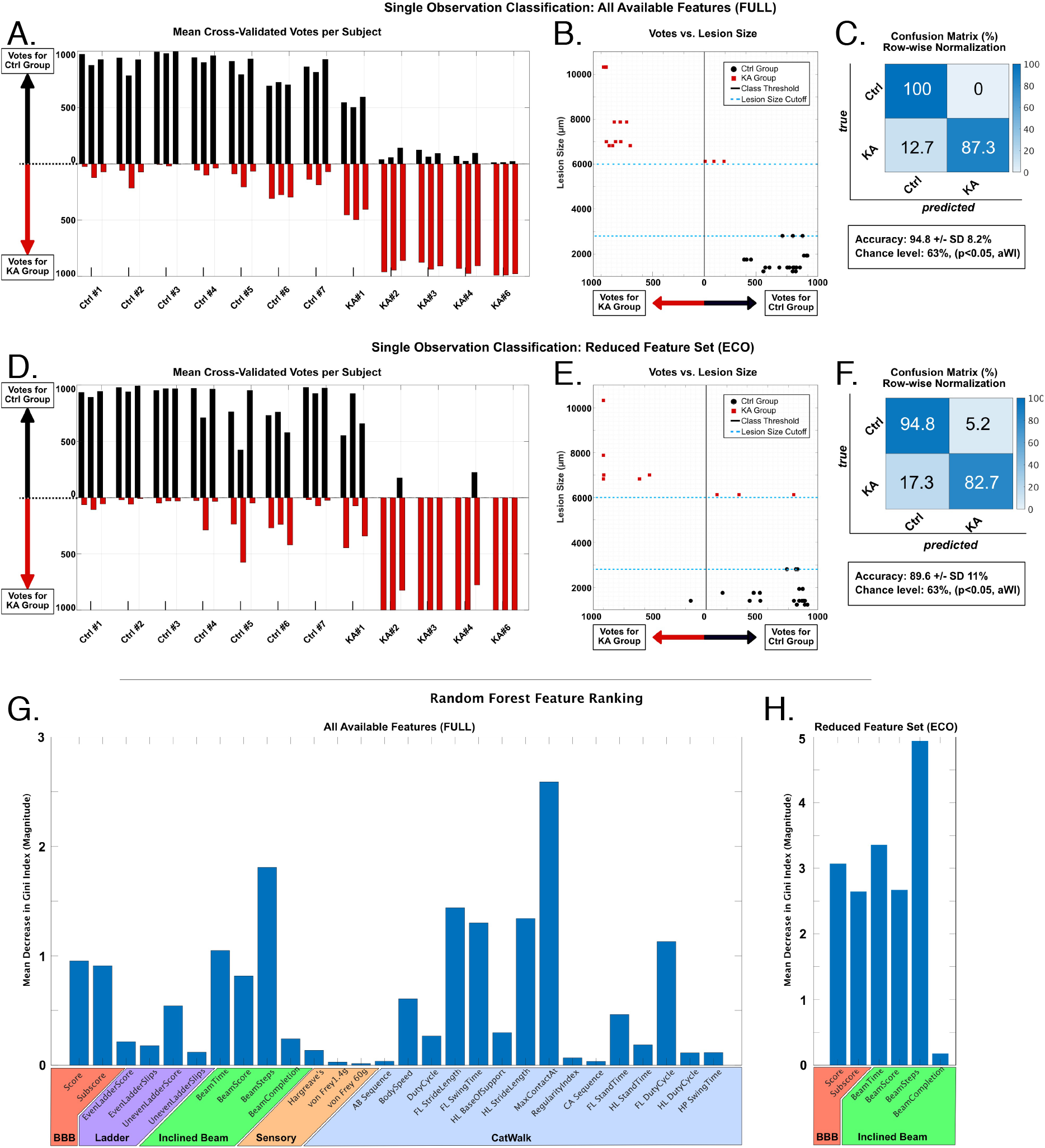
Behavioral classification of animals with the FULL and ECO models. ***A, D*** The left column shows the mean of the cross-validated votes (n = 1000) per observation per subject. For each animal, 3 observations were used (bar groups of three). Black bars indicate votes for the control group, red bars show votes for the KA group. Final class prediction was determined via majority vote. ***B, E*** The center column shows the relationship between votes (abscissa) and the post-hoc determined lesion size (ordinate). The class threshold is shown as horizontal line at point 0. The true class is shown in red for the KA group and in black for the control group. ***C, F*** The right column depicts the row-wise normalised confusion matrix for each classification approach as a percentage. The chance level was determined using an adjusted Wald Interval (aWI) at 63.0% (p<0.05). ***G, H*** Random Forest feature ranking according to the mean decrease in Gini Index for both FULL (**G**) and ECO (**H**) model. The higher the mean decrease, the more important the feature is considered to the classification model. N = 7 control and n = 5 KA animals.

As this information is important when applying treatments, we strove to develop a sensitive classification model that relied upon quick functional evaluation. A fast early classification approach used a reduced feature set (ECO) focusing only on the BBB and inclined beam behavioral tests and reached a mean accuracy of 89.6 ± SD 11%. The confusion matrix shows a misclassification rate of 5.2% (Type 1 error) for all control observations and 17.3% for all KA observations (Type 2 error) (**Fig. 9*F***). This is also reflected in the mean of the cross-validated votes (aggregated decisions of the ensembled decision trees) per subject: 11 out of 12 animals were also appropriately classified (**Fig. 9*D,E***). This Type 2 error seen in both models is understandable given the location and lesion extent of this animal (KA#1) compared to the rest of the animals, confirming our previous observations.

To validate the endpoint lesion size in the rostro-caudal axis as an exclusion criteria, we correlated the votes per trial with the corresponding animal lesion sizes and found significant correlations (**Fig. 9*B,E***: FULL model r = −0.7395 and p <0.007, ECO model r = −0.8916 and p <0.0001, Spearman’s correlation). The blue dotted lines indicate histologically determined cutoffs for the control and KA lesion sizes (2800µm and 6000µm, respectively) and further shows that the misclassified animal (KA#1) is the closest to this cutoff (**Fig. 9*B,E***). Together, these findings highlight the sensitivity and accuracy of the FULL and ECO models.

To determine which behavioral tests were most predictive and important for correct classification, a full ranking of the behavioral parameters was performed for both models and presented as the mean decrease of the Gini Index; the greater the decrease, the more discriminative information the behavioral test provides. For the FULL classification model, the highest rankings are achieved by the selected CatWalk parameters, however, all parameters obtained from the BBB and the inclined beam also showed consistent high rankings (**Fig. 9*G***). For the ECO model, rankings were all above 2.5 except for inclined beam completion (**Fig. 9*H***). Using these two models, combined behavioral performance and potential recovery after injury can be compared and evaluated both during the ongoing experiment with the ECO model, as well as post time-intensive analysis of all behavioral tests with the FULL model.

## Discussion

Targeted KA lesions to the intermediate gray matter (laminae V-VII) at spinal levels L2-L4 induced profound behavioral deficits related to locomotion. Damage to this area, which regulates lower motoneurons, leads not only to gross motor deficits (BBB score), but rhythmic and skilled walking (even and uneven horizontal ladders), coordination (BBB subscore), balance (inclined beam) and gait deficits (CatWalk), as well. We have shown that this lesion is specific to the intermediate gray matter in that the dorsal horn (laminae I-IV) remained relatively undisturbed (no significant KA-induced sensory deficits after two weeks) and we did not find correlations between motoneuron loss in the ventral horn or white matter damage and behavior. Neuronal loss in laminae V-VII correlated with coordination and balance deficits and provided further insight into the functional role of the IN populations spanning the spinal L2-L4 region. After three months, KA rats did not regain any lost sensorimotor function. This work indicates: one, the functional deficits of such an injury in this region; two, such an injury cannot be naturally compensated for; and three, that additional therapies must be considered for functional recovery.

### KA lesion targets local and propriospinal INs in laminae V-VII and are crucial for locomotion/coordination

In laminae V-VII of the intermediate gray matter reside both local SpINs and propriospinal INs which have been shown to play various regulatory roles in locomotion. Previous studies have found that treadmill walking activated spinal levels T13-L6 in adult rats, with significantly more cFos+ neurons in laminae IV, V and VII in trained versus untrained rats (Ahn et al., 2006). Furthermore, activity profiling of cFos+ SpINs in the mouse lumbar spinal cord following rotarod training revealed that many SpINs specific to motor activity (ie not active during baseline or painful formalin stimulation) are located in these laminae, including Gad2+ cells, NF1b+ cells and V2a neurons (Sathyamurthy et al., 2018). Therefore, it was likely this region would be integral to locomotion, however it remained to be seen what type of deficits would be displayed when the premotor excitatory/inhibitory IN ratio is shifted. We have found that collective damage to this area creates significant and lasting locomotor deficits that particularly affect coordination.

KA is an excitotoxin that has some selectivity for INs over motoneurons (Kwak & Nakamura, 1995; Urca & Urca, 1990). We have confirmed that the region of laminae V-VII is indeed successfully damaged with our lesion, however this does not exclude the possibility of some additional damage to surrounding lamina VIII. We confirmed that motoneurons and percent white matter did not correlate with behavioral deficits nor were significant sensory differences found between the control and KA groups. Neuronal damage to the dorsal horn can evoke pain-related behavior in mice (Yezierski et al., 1998) and animals with pain in their hindlimbs display an altered gait including in duty cycle and stride length parameters (Deshpande et al., 2021; Pitzer et al., 2016). The fact that there are no significant differences in mechanical or thermal sensitivity, suggests that KA did not affect the dorsal horn and impact our gait-related behavioral findings. These findings, therefore, indicate the specificity of our lesion and that damage to the SpINs in the intermediate gray matter which provide regulatory input to motoneurons resulted in the observed deficits.

### Lesioned premotor SpINs spanning L2-L4 and at least 6mm in length produce significant behavioral deficits

After behavioral analysis, we observed that the most predictive factors of a properly lesioned KA-animal were lesion size in the rostro-caudal axis spanning spinal L2-L4 and neuronal loss in laminae V-VII. We determined that KA-lesioned animals needed to have a minimum intermediate gray matter lesion size of at least 6mm in the rostro-caudal length to produce consistent behavioral deficits using our behavioral tests. In lesions shorter than 6mm, premotor circuitry in other spinal segments can compensate for the neuronal loss, resulting in very minor or no behavioral deficits (these animals were excluded from the analysis). In lesions greater than 6mm, we observed significant correlation with locomotor function after two weeks. The average lumbar cord (L1-L6) of the adult *Rattus norvegicus* spans approximately 18mm. Spinal levels L2-L4 measure approximately 10mm, of which each spinal segment spanned slightly over 3mm (Nicolopoulos-Stournaras & Iles, 1983). These findings suggest that the lesion size must span at least two spinal levels. For context regarding the premotor targeted circuitry, it should be noted that hindlimb muscles are innervated by motoneurons spanning several lumbar spinal levels (Liu et al., 2010; Puskar & Antal, 1997; Ronzano et al., 2021). The muscles vastus medialis, vastus lateralis, gracillis, tibialis anterior, biceps femoris muscles are innervated by motoneurons from at least two spinal levels in spinal L2-L4 (Mohan et al., 2015; Wenger et al., 2016). Therefore, the targeted regulatory circuitry of the motoneurons must also span a similar range. Although the lesions reported here are slightly longer than what has been previously published (3.4mm or 5.5mm), those previous lesions used 1.5µl of 5mM or 2.5mM KA per injection to produce larger gray matter lesions with more neuronal loss and also severely damaged dorsal horns (Magnuson et al., 1998).

While lesion size in the rostro-caudal axis plays an important role, it does not appear to be the only determining factor. For example, some animals have the same lesion extent, but varying behavioral deficits. We believe this may be due to the severity of the injury at given levels. The Random Forest classification and neuronal quantification highlight that this damage must include spinal level L2 to observe deficits in coordination and balance. This is most clearly demonstrated by animal KA #1, with a lesion size longer than 6mm (6125µm), exhibiting neuronal loss primarily in L3 and L4. The Random Forest classification of animal KA #1 produced a Type 2 error that lead to misclassification as a control due to its behavioral performance. All other animals with greater lesion sizes and neuronal loss were properly classified. This indicates that spinal L2 is integral to the observed behavioral loss. These findings support what has been previously published regarding circuitry critical to locomotion residing at this level in both the previously mentioned murine KA models and human models (Dimitrijevic et al., 1998; Hadi et al., 2000; Magnuson et al., 1999) and further illustrates the contribution of specifically the intermediate gray matter in locomotion.

### CatWalk gait analysis reveals deficits in rhythm and pattern generation

The CatWalk measures dynamic and static gait parameters. We initially focused on the pLDA score which draws on nine parameters that characterize contusion and dorsal hemisection SCI models. While both SCI models reside in the thoracic region, they target descending tracts which connect with lumbar premotor circuitry at varying degrees. Such differences were not categorized by the BBB gross hindlimb score but were highlighted by the pLDA score (Timotius et al., 2021). Our Gini Index results confirm that the CatWalk pLDA and subsequent parameters are highly predicative and that damage to gray matter in this particular region significantly alters gait.

CatWalk analysis revealed KA animals had rhythmic deficits (longer stand time and shorter swing time) however surprisingly these differences were primarily found in the forelimbs, not the hindlimbs. Furthermore, the Gini Index found several CatWalk forelimb parameters to be highly discriminant despite the lesion residing in the lumbar cord. Long propriospinal INs located in laminae VII-VIII connect the cervical and lumbar central pattern generator (CPG) networks (including in spinal levels C6-C8 and L1-L3) (Brockett et al., 2013; English et al., 1985; Laliberte et al., 2019; Pocratsky et al., 2020; Reed et al., 2006). This lesion model created neuronal loss in laminae V-VII, a region where several ascending propriospinal somas reside and descending propriospinal INs terminate (Flynn et al., 2011; Frigon, 2017). Therefore, while these changes could be a compensatory mechanism (Fouad et al., 2013), we hypothesize this may be due to propriospinal damage uncoupling the fore- and hindlimbs.

In addition to gait rhythm changes, we also saw significant differences in pattern generation. The regularity index measures correctly sequenced footsteps and is used to analyze recovery in mild to moderate injuries (Koopmans et al., 2005; Kuerzi et al., 2010; Shepard et al., 2021). As expected, KA-animals have a significantly lower regularity index, which indicates they have a lower paw-stepping quality.

Additionally, the step sequence change observed in KA animals could further be indicative of either long or short propriospinal damage. KA-lesioned animals have significantly higher CA step sequences, as opposed to the more commonly seen AB sequence which suggests interlimb coordination deficits (Baldwin et al., 2017) a characteristic also observed after long descending propriospinal ablation or stimulation (Ruder et al., 2016; Skinner et al., 1980). A previous study has shown that an increase in cruciate stepping was also observed after silencing descending spinal L2 short propriospinal INs which project to L5. This change in gait could therefore also be due to L2-L5 IN loss, as they are found in the intermediate gray matter (primarily laminae V-VIII) (Pocratsky et al., 2017), where our KA lesion also resides.

Together, these results show that KA-lesioned rats have deficits in gait rhythm and pattern parameters and further suggest propriospinal IN damage.

### Damage to the lumbar spinal enlargement is permanent and cannot be compensated

For this study, we were interested in determining how essential the intermediate gray matter of the lumbar cord is for locomotion and if redundant pathways or other neurons can compensate for premotor IN loss. Previous studies have shown that following SCI, propriospinal INs have the capacity to sprout and form new synaptic connections, with the potential to contribute to functional gains (Brommer et al., 2021; Courtine et al., 2008; Cowley et al., 2015; Shepard et al., 2021; Stelzner, 2008; Taccola et al., 2018; Zholudeva et al., 2018). The behavioral results of this study suggest that when lumbar spinal and propriospinal INs are lost, there are no other intrinsic mechanisms that can recover lost motor function. Hindlimb deficits level off after two weeks and remain stable even after three months. Strikingly, the two animals that did not have weight support after two weeks did not regain weight support at three months. Therefore, this model is well suited to investigate the efficacy of neurorestorative therapies after SCI in the lumbar spinal enlargements.

### Random Forest modeling determines classification prior to histological analysis

Experimental SCI models often have exclusion criteria to determine if the surgery was carried out properly, such as the displacement curve for contusion models (Cheriyan et al., 2014; Fournely et al., 2020; Sparrey et al., 2016). There is no such parameter available for this KA model, and lesion size and neuronal loss can only be determined post-mortem after time-intensive analysis. We aimed to create a mathematical model based on behavior that can be used to verify whether animals were properly lesioned. While Gini Index findings of the FULL model indicated that the select CatWalk parameters are the most predicative for proper classification, this apparatus is expensive and not easily available to all researchers. We therefore focused on the BBB and inclined beam behavioral tests for the ECO model, both of which use equipment that is more generally accessible, cost-effective and scoring can be completed quickly. By limiting the mathematical model to these two assessments, we slightly decrease reliability but keep a strong classification rate as observed in the ECO model (FULL model: 94.8% accuracy and ECO model: 89.6% accuracy). Our recommendation is for the ECO model to be used as an early exclusion criterion and for balancing group differences prior to intervention. However, this intervention would need to start after the two-week period when the behavioral data is collected and analyzed, therefore it lends itself more effectively to cellular replacement therapies. The FULL model provides a more sensitive behavioral evaluation than a single test alone and can be used post-experiment to analyze intervention effects.

### Ideas and Speculation

The results of this study highlight the role intermediate gray matter SpINs play in the lumbar enlargement. Furthermore, they do not seem to be compensated for by spontaneous remodeling. Previous studies have compared contusion and KA injuries in both the thoracic and lumbar cord and have shown that the lumbar gray matter plays a greater role in locomotion than the thoracic gray matter (Magnuson et al., 1998). Our findings show that damage to the lumbar intermediate gray matter creates lasting deficits beyond gross hindlimb function and particularly affects coordination. Therefore, while many SCI treatments focus on regeneration of host neurons, primarily of the white matter tracts, other treatments such as gray matter replacement therapies are required for injuries to the lumbar enlargement. This new SCI model with fully characterized behavioral deficits and sensitive neuronal quantification and classification models will help carefully evaluate the potential of a cellular replacement therapy and aid future SCI treatment options.

## Supporting information

Kuehn et al. Supplementary movie

Kuehn et al. Supplementary Table and Figures

## Footnotes

This research was supported by Olympia Morata Program Fellowship of the University of Heidelberg Faculty of Medicine awarded to R. Puttagunta as well as grants from the Deutsche Forschungsgemeinschaft (SFB 1158 A06, PU 425/4-3 and WE 2165/4-3) awarded to R.Puttagunta and N.Weidner. Behavioral CatWalk studies were supported by the Interdisciplinary Neuroscience Behavioral Core (INBC) at Heidelberg University. We gratefully acknowledge the data storage service SDS@hd supported by the Ministry of Science, Research and the Arts Baden-Württemberg (MWK). We would like to especially thank Dr. David Magnuson and Alice Shum-Siu for sharing their expertise and advice to establish the kainic acid lesion model. We would also like to thank Dr. Thomas Schackel for his help with surgical training.

### Abbreviations

CPG: central pattern generator
CSA: cross sectional area
dpi: days post-injury
INs: interneurons
KA: kainic acid
MIP: maximum intensity projection
ROIs: regions of interest
SCI: spinal cord injury
SD: standard deviation
SEM: standard error of the mean
SpINs: spinal interneurons

## References

Ahn, S. N., Guu, J. J., Tobin, A. J., Edgerton, V. R., & Tillakaratne, N. J. K. (2006). Use of c-fos to identify activity-dependent spinal neurons after stepping in intact adult rats. Spinal Cord, 44(9), 547–559. https://doi.org/10.1038/sj.sc.3101862

Baldwin, H. A., Koivula, P. P., Necarsulmer, J. C., Whitaker, K. W., & Harvey, B. K. (2017). Step Sequence Is a Critical Gait Parameter of Unilateral 6-OHDA Parkinson’s Rat Models. Cell Transplantation, 26(4), 659–667. https://doi.org/10.3727/096368916x693059

Basso, D. M., Beattie, M. S., & Bresnahan, J. C. (1995). A sensitive and reliable locomotor rating scale for open field testing in rats. J Neurotrauma, 12(1), 1–21. https://doi.org/10.1089/neu.1995.12.1

Berg, S., Kutra, D., Kroeger, T., Straehle, C. N., Kausler, B. X., Haubold, C., Schiegg, M., Ales, J., Beier, T., Rudy, M., Eren, K., Cervantes, J. I., Xu, B., Beuttenmueller, F., Wolny, A., Zhang, C., Koethe, U., Hamprecht, F. A., & Kreshuk, A. (2019). ilastik: interactive machine learning for (bio)image analysis. Nat Methods, 16(12), 1226–1232. https://doi.org/10.1038/s41592-019-0582-9

Breiman, L. (2001). Random forests. Machine Learning, 45(1), 5–32. https://doi.org/Doi10.1023/A:1010933404324

Brockett, E. G., Seenan, P. G., Bannatyne, B. A., & Maxwell, D. J. (2013). Ascending and descending propriospinal pathways between lumbar and cervical segments in the rat: evidence for a substantial ascending excitatory pathway. Neuroscience, 240, 83–97. https://doi.org/10.1016/j.neuroscience.2013.02.039

Brommer, B., He, M., Zhang, Z., Yang, Z., Page, J. C., Su, J., Zhang, Y., Zhu, J., Gouy, E., Tang, J., Williams, P., Dai, W., Wang, Q., Solinsky, R., Chen, B., & He, Z. (2021). Improving hindlimb locomotor function by Non-invasive AAV-mediated manipulations of propriospinal neurons in mice with complete spinal cord injury. Nat Commun, 12(1), 781. https://doi.org/10.1038/s41467-021-20980-4

Carter, R. J., Morton, J., & Dunnett, S. B. (2001). Motor coordination and balance in rodents. Curr Protoc Neurosci, Chapter 8, Unit 8 12. https://doi.org/10.1002/0471142301.ns0812s15

Cheriyan, T., Ryan, D. J., Weinreb, J. H., Cheriyan, J., Paul, J. C., Lafage, V., Kirsch, T., & Errico, T. J. (2014). Spinal cord injury models: a review. Spinal Cord, 52(8), 588–595. https://doi.org/10.1038/sc.2014.91

Courtine, G., Song, B., Roy, R. R., Zhong, H., Herrmann, J. E., Ao, Y., Qi, J., Edgerton, V. R., & Sofroniew, M. V. (2008). Recovery of supraspinal control of stepping via indirect propriospinal relay connections after spinal cord injury. Nat Med, 14(1), 69–74. https://doi.org/10.1038/nm1682

Cowley, K. C., MacNeil, B. J., Chopek, J. W., Sutherland, S., & Schmidt, B. J. (2015). Neurochemical excitation of thoracic propriospinal neurons improves hindlimb stepping in adult rats with spinal cord lesions. Exp Neurol, 264, 174–187. https://doi.org/10.1016/j.expneurol.2014.12.006

Dai, X., Noga, B. R., Douglas, J. R., & Jordan, L. M. (2005). Localization of spinal neurons activated during locomotion using the c-fos immunohistochemical method. J Neurophysiol, 93(6), 3442–3452. https://doi.org/10.1152/jn.00578.2004

Deshpande, D., Agarwal, N., Fleming, T., Gaveriaux-Ruff, C., Klose, C. S. N., Tappe-Theodor, A., Kuner, R., & Nawroth, P. (2021). Loss of POMC-mediated antinociception contributes to painful diabetic neuropathy. Nat Commun, 12(1), 426. https://doi.org/10.1038/s41467-020-20677-0

Dimitrijevic, M. R., Gerasimenko, Y., & Pinter, M. M. (1998). Evidence for a spinal central pattern generator in humans. Ann N Y Acad Sci, 860, 360–376. https://doi.org/10.1111/j.1749-6632.1998.tb09062.x

English, A. W., Tigges, J., & Lennard, P. R. (1985). Anatomical organization of long ascending propriospinal neurons in the cat spinal cord. J Comp Neurol, 240(4), 349–358. https://doi.org/10.1002/cne.902400403

Flynn, J. R., Graham, B. A., Galea, M. P., & Callister, R. J. (2011). The role of propriospinal interneurons in recovery from spinal cord injury. Neuropharmacology, 60(5), 809–822. https://doi.org/10.1016/j.neuropharm.2011.01.016

Fouad, K., Hurd, C., & Magnuson, D. S. (2013). Functional testing in animal models of spinal cord injury: not as straight forward as one would think. Front Integr Neurosci, 7, 85. https://doi.org/10.3389/fnint.2013.00085

Fournely, M., Petit, Y., Wagnac, E., Evin, M., & Arnoux, P. J. (2020). Effect of experimental, morphological and mechanical factors on the murine spinal cord subjected to transverse contusion: A finite element study. PLoS One, 15(5), e0232975. https://doi.org/10.1371/journal.pone.0232975

Frigon, A. (2017). The neural control of interlimb coordination during mammalian locomotion. J Neurophysiol, 117(6), 2224–2241. https://doi.org/10.1152/jn.00978.2016

Gosgnach, S., Lanuza, G. M., Butt, S. J., Saueressig, H., Zhang, Y., Velasquez, T., Riethmacher, D., Callaway, E. M., Kiehn, O., & Goulding, M. (2006). V1 spinal neurons regulate the speed of vertebrate locomotor outputs. Nature, 440(7081), 215–219. https://doi.org/10.1038/nature04545

Guertin, P. A. (2009). The mammalian central pattern generator for locomotion. Brain Res Rev, 62(1), 45–56. https://doi.org/10.1016/j.brainresrev.2009.08.002

Hadi, B., Zhang, Y. P., Burke, D. A., Shields, C. B., & Magnuson, D. S. (2000). Lasting paraplegia caused by loss of lumbar spinal cord interneurons in rats: no direct correlation with motor neuron loss. J Neurosurg, 93(2 Suppl), 266–275. https://doi.org/10.3171/spi.2000.93.2.0266

Hayashi, M., Hinckley, C. A., Driscoll, S. P., Moore, N. J., Levine, A. J., Hilde, K. L., Sharma, K., & Pfaff, S. L. (2018). Graded Arrays of Spinal and Supraspinal V2a Interneuron Subtypes Underlie Forelimb and Hindlimb Motor Control. Neuron, 97(4), 869–884 e865. https://doi.org/10.1016/j.neuron.2018.01.023

Kerchner, G. A., Wang, G. D., Qiu, C. S., Huettner, J. E., & Zhuo, M. (2001). Direct presynaptic regulation of GABA/glycine release by kainate receptors in the dorsal horn: an ionotropic mechanism. Neuron, 32(3), 477–488. https://doi.org/10.1016/s0896-6273(01)00479-2

Koch, S. C., Del Barrio, M. G., Dalet, A., Gatto, G., Gunther, T., Zhang, J., Seidler, B., Saur, D., Schule, R., & Goulding, M. (2017). RORbeta Spinal Interneurons Gate Sensory Transmission during Locomotion to Secure a Fluid Walking Gait. Neuron, 96(6), 1419–1431 e1415. https://doi.org/10.1016/j.neuron.2017.11.011

Koopmans, G. C., Deumens, R., Honig, W. M., Hamers, F. P., Steinbusch, H. W., & Joosten, E. A. (2005). The assessment of locomotor function in spinal cord injured rats: the importance of objective analysis of coordination. J Neurotrauma, 22(2), 214–225. https://doi.org/10.1089/neu.2005.22.214

Kuerzi, J., Brown, E. H., Shum-Siu, A., Siu, A., Burke, D., Morehouse, J., Smith, R. R., & Magnuson, D. S. (2010). Task-specificity vs. ceiling effect: step-training in shallow water after spinal cord injury. Exp Neurol, 224(1), 178–187. https://doi.org/10.1016/j.expneurol.2010.03.008

Kwak, S., & Nakamura, R. (1995). Selective degeneration of inhibitory interneurons in the rat spinal cord induced by intrathecal infusion of acromelic acid. Brain Res, 702(1-2), 61–71. https://doi.org/10.1016/0006-8993(95)01000-6

Laliberte, A. M., Goltash, S., Lalonde, N. R., & Bui, T. V. (2019). Propriospinal Neurons: Essential Elements of Locomotor Control in the Intact and Possibly the Injured Spinal Cord. Front Cell Neurosci, 13, 512. https://doi.org/10.3389/fncel.2019.00512

Lanuza, G. M., Gosgnach, S., Pierani, A., Jessell, T. M., & Goulding, M. (2004). Genetic identification of spinal interneurons that coordinate left-right locomotor activity necessary for walking movements. Neuron, 42(3), 375–386. https://doi.org/10.1016/s0896-6273(04)00249-1

Liu, T. T., Bannatyne, B. A., & Maxwell, D. J. (2010). Organization and neurochemical properties of intersegmental interneurons in the lumbar enlargement of the adult rat. Neuroscience, 171(2), 461–484. https://doi.org/10.1016/j.neuroscience.2010.09.012

Magnuson, D., Trinder, T., Zhang, Y. P., Burke, D., Morassutti, D., & Shields, C. (1998). Comparing Deficits Following Excitotoxic and Contusion Injuries in the Thoracic and Lumbar Spinal Cord of the Adult Rat. In (Vol. 156, pp. 191–204): Experimental Neurology.

Magnuson, D. S., Lovett, R., Coffee, C., Gray, R., Han, Y., Zhang, Y. P., & Burke, D. A. (2005). Functional consequences of lumbar spinal cord contusion injuries in the adult rat. J Neurotrauma, 22(5), 529–543. https://doi.org/10.1089/neu.2005.22.529

Magnuson, D. S., Trinder, T. C., Zhang, Y. P., Burke, D., Morassutti, D. J., & Shields, C. B. (1999). Comparing deficits following excitotoxic and contusion injuries in the thoracic and lumbar spinal cord of the adult rat. Exp Neurol, 156(1), 191–204. https://doi.org/10.1006/exnr.1999.7016

Martins, L. A., Schiavo, A., Xavier, L. L., & Mestriner, R. G. (2022). The Foot Fault Scoring System to Assess Skilled Walking in Rodents: A Reliability Study. Front Behav Neurosci, 16, 892010. https://doi.org/10.3389/fnbeh.2022.892010

Metz, G. A., & Whishaw, I. Q. (2009). The ladder rung walking task: a scoring system and its practical application. J Vis Exp(28). https://doi.org/10.3791/1204

Mohan, R., Tosolini, A. P., & Morris, R. (2015). Segmental Distribution of the Motor Neuron Columns That Supply the Rat Hindlimb: A Muscle/Motor Neuron Tract-Tracing Analysis Targeting the Motor End Plates. Neuroscience, 307, 98–108. https://doi.org/10.1016/j.neuroscience.2015.08.030

Müller-Putz, G. R., Scherer, R., Brunner, C., Leeb, R., & Pfurtscheller, G. (2008). Better than random? A closer look on BCI results. International Journal of Bioelectromagnetism, 10(1), 4.

Nicolopoulos-Stournaras, S., & Iles, J. F. (1983). Motor neuron columns in the lumbar spinal cord of the rat. J Comp Neurol, 217(1), 75–85. https://doi.org/10.1002/cne.902170107

Percie du Sert, N., Hurst, V., Ahluwalia, A., Alam, S., Avey, M. T., Baker, M., Browne, W. J., Clark, A., Cuthill, I. C., Dirnagl, U., Emerson, M., Garner, P., Holgate, S. T., Howells, D. W., Karp, N. A., Lazic, S. E., Lidster, K., MacCallum, C. J., Macleod, M., Wurbel, H. (2020). The ARRIVE guidelines 2.0: Updated guidelines for reporting animal research. PLoS Biol, 18(7), e3000410. https://doi.org/10.1371/journal.pbio.3000410

Pitzer, C., Kuner, R., & Tappe-Theodor, A. (2016). Voluntary and evoked behavioral correlates in neuropathic pain states under different social housing conditions. Molecular Pain, 12. https://doi.org/Artn174480691665663510.1177/1744806916656635

Pocratsky, A. M., Burke, D. A., Morehouse, J. R., Beare, J. E., Riegler, A. S., Tsoulfas, P., States, G. J. R., Whittemore, S. R., & Magnuson, D. S. K. (2017). Reversible silencing of lumbar spinal interneurons unmasks a task-specific network for securing hindlimb alternation. Nature Communications, 8. https://doi.org/ARTN196310.1038/s41467-017-02033-x

Pocratsky, A. M., Shepard, C. T., Morehouse, J. R., Burke, D. A., Riegler, A. S., Hardin, J. T., Beare, J. E., Hainline, C., States, G. J., Brown, B. L., Whittemore, S. R., & Magnuson, D. S. (2020). Long ascending propriospinal neurons provide flexible, context-specific control of interlimb coordination. Elife, 9. https://doi.org/10.7554/eLife.53565

Preibisch, S., Saalfeld, S., & Tomancak, P. (2009). Globally optimal stitching of tiled 3D microscopic image acquisitions. Bioinformatics, 25(11), 1463–1465. https://doi.org/10.1093/bioinformatics/btp184

Puskar, Z., & Antal, M. (1997). Localization of last-order premotor interneurons in the lumbar spinal cord of rats. J Comp Neurol, 389(3), 377–389. https://doi.org/10.1002/(sici)1096-9861(19971222)389:3<377::aid-cne2>3.0.co;2-y

Reed, W. R., Shum-Siu, A., Onifer, S. M., & Magnuson, D. S. (2006). Inter-enlargement pathways in the ventrolateral funiculus of the adult rat spinal cord. Neuroscience, 142(4), 1195–1207. https://doi.org/10.1016/j.neuroscience.2006.07.017

Rodriguez-Moreno, A., Lopez-Garcia, J. C., & Lerma, J. (2000). Two populations of kainate receptors with separate signaling mechanisms in hippocampal interneurons. Proc Natl Acad Sci U S A, 97(3), 1293–1298. https://doi.org/10.1073/pnas.97.3.1293

Ronzano, R., Lancelin, C., Bhumbra, G. S., Brownstone, R. M., & Beato, M. (2021). Proximal and distal spinal neurons innervating multiple synergist and antagonist motor pools. Elife, 10. https://doi.org/10.7554/eLife.70858

Ruder, L., Takeoka, A., & Arber, S. (2016). Long-Distance Descending Spinal Neurons Ensure Quadrupedal Locomotor Stability. Neuron, 92(5), 1063–1078. https://doi.org/10.1016/j.neuron.2016.10.032

Sathyamurthy, A., Johnson, K. R., Matson, K. J. E., Dobrott, C. I., Li, L., Ryba, A. R., Bergman, T. B., Kelly, M. C., Kelley, M. W., & Levine, A. J. (2018). Massively Parallel Single Nucleus Transcriptional Profiling Defines Spinal Cord Neurons and Their Activity during Behavior. Cell Rep, 22(8), 2216–2225. https://doi.org/10.1016/j.celrep.2018.02.003

Schindelin, J., Arganda-Carreras, I., Frise, E., Kaynig, V., Longair, M., Pietzsch, T., Preibisch, S., Rueden, C., Saalfeld, S., Schmid, B., Tinevez, J. Y., White, D. J., Hartenstein, V., Eliceiri, K., Tomancak, P., & Cardona, A. (2012). Fiji: an open-source platform for biological-image analysis. Nat Methods, 9(7), 676–682. https://doi.org/10.1038/nmeth.2019

Shepard, C. T., Pocratsky, A. M., Brown, B. L., Van Rijswijck, M. A., Zalla, R. M., Burke, D. A., Morehouse, J. R., Riegler, A. S., Whittemore, S. R., & Magnuson, D. S. (2021). Silencing long ascending propriospinal neurons after spinal cord injury improves hindlimb stepping in the adult rat. Elife, 10. https://doi.org/10.7554/eLife.70058

Skinner, R. D., Adams, R. J., & Remmel, R. S. (1980). Responses of Long Descending Propriospinal Neurons to Natural and Electrical Types of Stimuli in Cat. Brain Research, 196(2), 387–403. https://doi.org/Doi10.1016/0006-8993(80)90403-5

Sliwinski, C., Nees, T. A., Puttagunta, R., Weidner, N., & Blesch, A. (2018). Sensorimotor Activity Partially Ameliorates Pain and Reduces Nociceptive Fiber Density in the Chronically Injured Spinal Cord. J Neurotrauma, 35(18), 2222–2238. https://doi.org/10.1089/neu.2017.5431

Smith, A. J. (2020). Guidelines for planning and conducting high-quality research and testing on animals. Lab Anim Res, 36, 21. https://doi.org/10.1186/s42826-020-00054-0

Sparrey, C. J., Salegio, E. A., Camisa, W., Tam, H., Beattie, M. S., & Bresnahan, J. C. (2016). Mechanical Design and Analysis of a Unilateral Cervical Spinal Cord Contusion Injury Model in Non-Human Primates. J Neurotrauma, 33(12), 1136–1149. https://doi.org/10.1089/neu.2015.3974

Stelzner, D. J. (2008). Short-circuit recovery from spinal injury. Nat Med, 14(1), 19–20. https://doi.org/10.1038/nm0108-19

Sternfeld, M. J., Hinckley, C. A., Moore, N. J., Pankratz, M. T., Hilde, K. L., Driscoll, S. P., Hayashi, M., Amin, N. D., Bonanomi, D., Gifford, W. D., Sharma, K., Goulding, M., & Pfaff, S. L. (2017). Speed and segmentation control mechanisms characterized in rhythmically-active circuits created from spinal neurons produced from genetically-tagged embryonic stem cells. Elife, 6. https://doi.org/10.7554/eLife.21540

Stringer, C., Wang, T., Michaelos, M., & Pachitariu, M. (2021). Cellpose: a generalist algorithm for cellular segmentation. Nat Methods, 18(1), 100–106. https://doi.org/10.1038/s41592-020-01018-x

Taccola, G., Sayenko, D., Gad, P., Gerasimenko, Y., & Edgerton, V. R. (2018). And yet it moves: Recovery of volitional control after spinal cord injury. Prog Neurobiol, 160, 64–81. https://doi.org/10.1016/j.pneurobio.2017.10.004

Takahashi, Y., Chiba, T., Kurokawa, M., & Aoki, Y. (2003). Dermatomes and the central organization of dermatomes and body surface regions in the spinal cord dorsal horn in rats. Journal of Comparative Neurology, 462(1), 29–41. https://doi.org/10.1002/cne.10669

Timotius, I. K., Bieler, L., Couillard-Despres, S., Sandner, B., Garcia-Ovejero, D., Labombarda, F., Estrada, V., Müller, H. W., Winkler, J., Klucken, J., Eskofier, B., Weidner, N., & Puttagunta, R. (2021). Combination of Defined CatWalk Gait Parameters for Predictive Locomotion Recovery in Experimental Spinal Cord Injury Rat Models. eNeuro, 8(2). https://doi.org/10.1523/ENEURO.0497-20.2021

Urca, G., & Urca, R. (1990). Neurotoxic effects of excitatory amino acids in the mouse spinal cord: quisqualate and kainate but not N-methyl-D-aspartate induce permanent neural damage. Brain Res, 529(1-2), 7–15. https://doi.org/10.1016/0006-8993(90)90805-l

Vierck, C. J., King, C. D., Berens, S. A., & Yezierski, R. P. (2013). Excitotoxic injury to thoracolumbar gray matter alters sympathetic activation and thermal pain sensitivity. Exp Brain Res, 231(1), 19–26. https://doi.org/10.1007/s00221-013-3666-2

Watson, C., Paxinos, G., & Kayaliuglu, G. (2009). The Spinal Cord. Elsevier Academic Press.

Wen, J., Sun, D., Tan, J., & Young, W. (2015). A consistent, quantifiable, and graded rat lumbosacral spinal cord injury model. J Neurotrauma, 32(12), 875–892. https://doi.org/10.1089/neu.2013.3321

Wenger, N., Moraud, E. M., Gandar, J., Musienko, P., Capogrosso, M., Baud, L., Le Goff, C. G., Barraud, Q., Pavlova, N., Dominici, N., Minev, I. R., Asboth, L., Hirsch, A., Duis, S., Kreider, J., Mortera, A., Haverbeck, O., Kraus, S., Schmitz, F., … Courtine, G. (2016). Spatiotemporal neuromodulation therapies engaging muscle synergies improve motor control after spinal cord injury. Nat Med, 22(2), 138–145. https://doi.org/10.1038/nm.4025

Yezierski, P. R., Liu, S., Ruenes, L. G., Kajander, J. K., & Brewer, L. K. (1998). Excitotoxic spinal cord injury: behavioral and morphological characteristics of a central pain model. Pain, 75(1), 141–155. https://doi.org/10.1016/S0304-3959(97)00216-9

Zhang, J., Lanuza, G. M., Britz, O., Wang, Z., Siembab, V. C., Zhang, Y., Velasquez, T., Alvarez, F. J., Frank, E., & Goulding, M. (2014). V1 and v2b interneurons secure the alternating flexor-extensor motor activity mice require for limbed locomotion. Neuron, 82(1), 138–150. https://doi.org/10.1016/j.neuron.2014.02.013

Zholudeva, L. V., Abraira, V. E., Satkunendrarajah, K., McDevitt, T. C., Goulding, M. D., Magnuson, D. S. K., & Lane, M. A. (2021). Spinal Interneurons as Gatekeepers to Neuroplasticity after Injury or Disease. J Neurosci, 41(5), 845–854. https://doi.org/10.1523/JNEUROSCI.1654-20.2020

Zholudeva, L. V., Qiang, L., Marchenko, V., Dougherty, K. J., Sakiyama-Elbert, S. E., & Lane, M. A. (2018). The Neuroplastic and Therapeutic Potential of Spinal Interneurons in the Injured Spinal Cord. Trends Neurosci, 41(9), 625–639. https://doi.org/10.1016/j.tins.2018.06.004

