## Supplementary material for "Intermediate Gray Matter Interneurons in the Lumbar Spinal Cord Play a Critical and Necessary Role in Coordinated Locomotion": Kuehn et al. Supplementary Table and Figures

**Supplementary Table 1.** Feature extraction and observation generation for the FULL model.

| # | Feature Type | # Repetitions | Method |
|---|---|---|---|
| 1 | 'BBBScore' | 1 | Duplicate for each observation per animal |
| 2 | 'BBBSubscore' | 1 | Duplicate for each observation per animal |
| 3 | 'EvenLadderScore' | [5 1] | First 3 repetitions |
| 4 | 'EvenLadderSlips' | [5 1] | First 3 repetitions |
| 5 | 'UnevenLadderScore' | [5 1] | First 3 repetitions |
| 6 | 'UnevenLadderSlips' | [5 1] | First 3 repetitions |
| 7 | 'ICBeamTime' | [3 1] | First 3 repetitions |
| 8 | 'ICBeamScore' | [3 1] | First 3 repetitions |
| 9 | 'ICBeamSteps' | [3 1] | First 3 repetitions |
| 10 | 'ICBeamCompletions' | [3 1] | First 3 repetitions |
| 11 | 'Hargreaves' | [4 1] | First 3 repetitions |
| 12 | 'Frey1_4' | [5 1] | First 3 repetitions |
| 13 | 'Frey60' | [5 1] | First 3 repetitions |
| 14 | 'CWABSeq' | [5 1] | First 3 repetitions |
| 15 | 'CWBodySpeed' | [5 1] | First 3 repetitions |
| 16 | 'CWDutyCycle' | [5 1] | First 3 repetitions |
| 17 | 'CWFLStrideLength' | [5 1] | First 3 repetitions |
| 18 | 'CWFLSwingTime' | [5 1] | First 3 repetitions |
| 19 | 'CWHLBaseOfSupport' | [5 1] | First 3 repetitions |
| 20 | 'CatwalkHLStrideLength' | [5 1] | First 3 repetitions |
| 21 | 'CWMaxContactAt' | [5 1] | First 3 repetitions |
| 22 | 'CWRegIdx' | [5 1] | First 3 repetitions |
| 23 | 'CWCASequence' | [5 1] | First 3 repetitions |
| 24 | 'CWFLStandTime' | [5 1] | First 3 repetitions |
| 25 | 'CWHLStandTime' | [5 1] | First 3 repetitions |
| 26 | 'CWFLDutyCycle' | [5 1] | First 3 repetitions |
| 27 | 'CWHLDutyCycle' | [5 1] | First 3 repetitions |
| 28 | 'CWHLSwingTime' | [5 1] | First 3 repetitions |

**Supplementary Table 2.** Feature extraction and observation generation for the ECO model.

| # | Feature Type | # Repetitions | Method |
|---|---|---|---|
| 1 | 'BBBScore' | 1 | Duplicate for each observation per animal |
| 2 | 'BBBSubscore' | 1 | Duplicate for each observation per animal |
| 3 | 'ICBeamTime' | [3 1] | First 3 repetitions |
| 4 | 'ICBeamScore' | [3 1] | First 3 repetitions |
| 5 | 'ICBeamSteps' | [3 1] | First 3 repetitions |
| 6 | 'ICBeamCompletions' | [3 1] | First 3 repetitions |

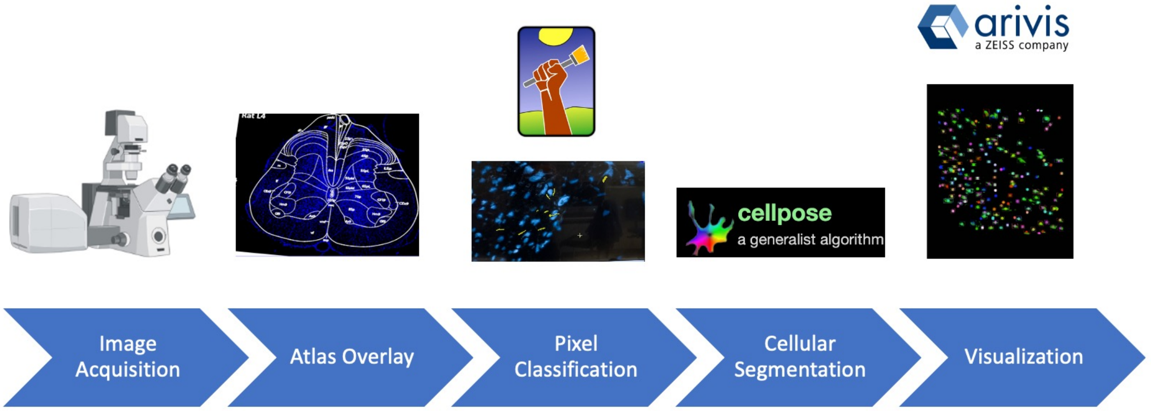


**Supplementary Figure 1. Image acquisition and analysis workflow designed for neuronal quantification.** First, Z-stack tiles of coronal slices stained with a NeuN antibody were acquired using a confocal XT1000 microscope with the 10x magnification objective. The tiles were stitched in ImageJ/Fiji and a spinal cord atlas overlay was registered over the maximum intensity projection using the BUnwarpJ ImageJ/Fiji plugin. Once the correct spinal levels and ROIs were determined (laminas V-VII), the ilastik pixel classification workflow was trained, and the output foreground probability map used as an input in cellpose to 3D segment the nuclei (cellpose, nuclei pretrained model). The 3D labeled images were visualized in arivis Vision4D and segmentation mistakes were manually corrected. The neuronal counts were normalized by ROI volume.

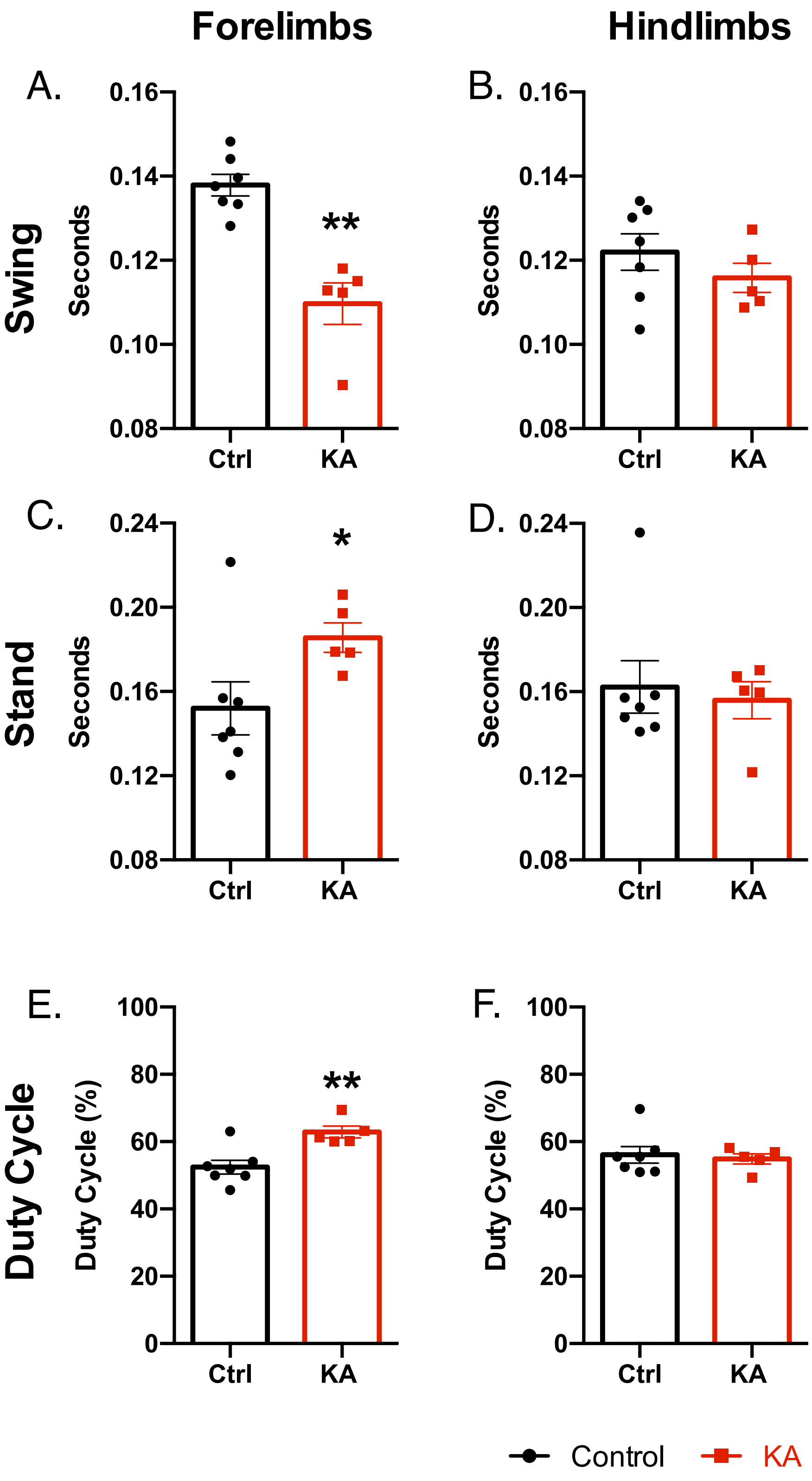


**Supplementary Figure 2. Rhythmic component in gait is significantly affected in KA-injured animals two weeks post-injury.** ***A-F*** Average swing time, stand time and duty cycle were significantly different for the forelimbs but not for the hindlimbs (Welch's unpaired t-test ***A,*** p = 0.022; ***B,*** p = 0.2939; ***C,*** p = 0.0441; ***D,*** p = 0.6858; ***E,*** p = 0.0031; ***F,*** = 0.6807). N = 7 control and n = 5 KA animals; * $p \leq 0.05$, ** $p \leq 0.01$.

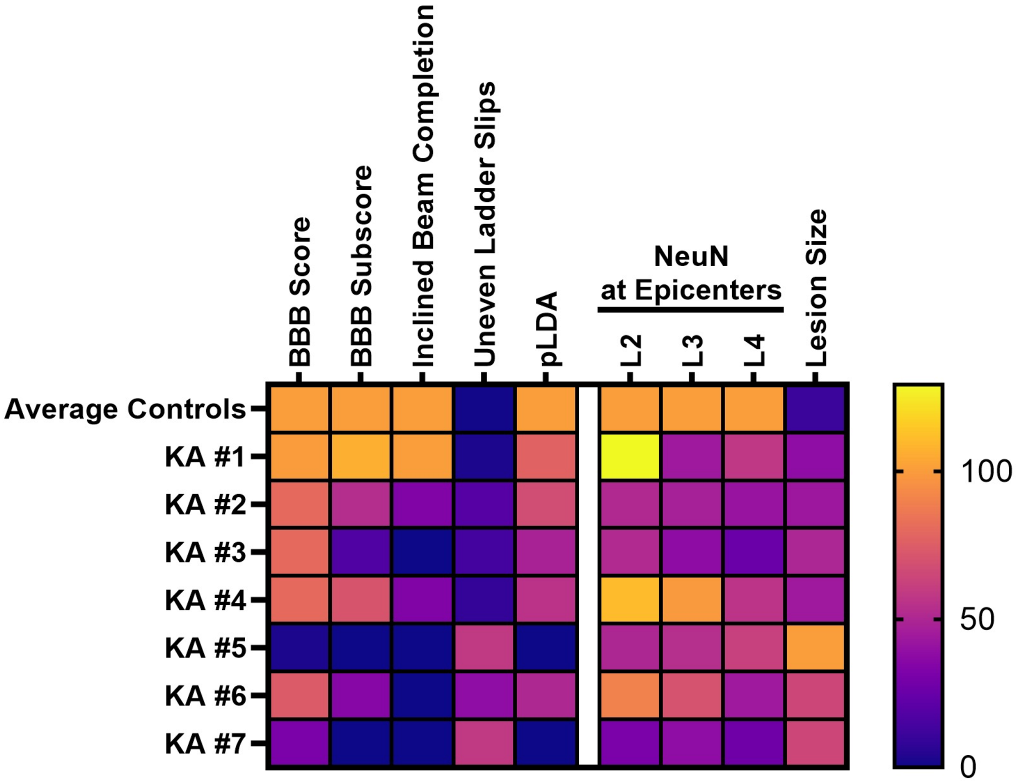


**Supplementary Figure 3. Representative heatmap comparing control and KA NeuN-positive cells in laminae V-VII in spinal levels L2-L4 to behavioral performance.** All neuronal values and behavioral performances are normalized to the average controls; the lesion size is normalized to the largest lesion extent (KA #5). All values are shown as a percentage (0% in violet, 100% in orange, above 100% in yellow).

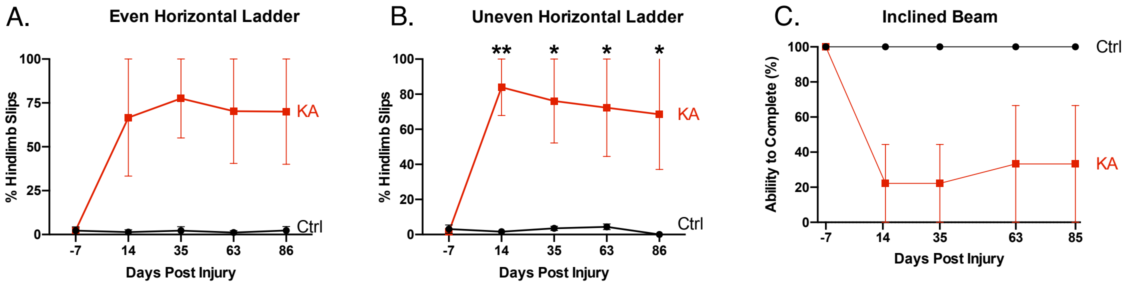


**Supplementary Figure 4**. **Coordination deficits in KA animals remain after three months.** ***A,*** Percent hindlimb slips on the even horizontal ladder is compared between the two groups (2-way ANOVA with Sidak's post hoc test, group p = 0.0767). ***B,*** Percent hindlimb slips on the uneven horizontal ladder is compared between the two groups (2-way ANOVA with Sidak's post hoc test, group p = 0.0442). ***C,*** Ability to complete the inclined beam over three months is compared between the two groups (2-way ANOVA with Sidak's post hoc test, group p = 0.1375). N = 3 animals per group; * p ≤ 0.05, ** p ≤ 0.01.
